# *Geobacter sulfurreducens* inner membrane cytochrome CbcBA controls electron transfer and growth yield near the energetic limit of respiration

**DOI:** 10.1101/2021.04.15.440034

**Authors:** Komal Joshi, Chi Ho Chan, Daniel R. Bond

## Abstract

*Geobacter sulfurreducens* utilizes extracellular electron acceptors such as Mn(IV), Fe(III), syntrophic partners, and electrodes that vary from +0.4 to −0.3 V *vs.* Standard Hydrogen Electrode (SHE), representing a potential energy span that should require a highly branched electron transfer chain. Here we describe CbcBA, a *bc*-type cytochrome essential near the thermodynamic limit of respiration when acetate is the electron donor. Mutants lacking *cbcBA* ceased Fe(III) reduction at −0.21 V *vs.* SHE, could not transfer electrons to electrodes between −0.21 and −0.28 V, and could not reduce the final 10% – 35% of Fe(III) minerals. As redox potential decreased during Fe(III) reduction, *cbcBA* was induced with the aid of the regulator BccR to become one of the most highly expressed genes in *G. sulfurreducens*. Growth yield (CFU/mM Fe(II)) was 112% of WT in Δ*cbcBA*, and deletion of *cbcL* (a different *bc*-cytochrome essential near −0.15 V) in Δ*cbcBA* increased yield to 220%. Together with ImcH, which is required at high redox potentials, CbcBA represents a third cytoplasmic membrane oxidoreductase in *G. sulfurreducens*. This expanding list shows how these important metal-reducing bacteria may constantly sense redox potential to adjust growth efficiency in changing environments.

## Introduction

Life generates cellular energy by linking electron donor oxidation to acceptor reduction. Each electron source and sink has an inherent affinity for electrons, or redox potential, which defines the maximum amount of energy available in such coupled reactions. For example, the difference in midpoint potentials of NO_3−_ and Fe(III) is more than half a volt, which is enough to generate an additional ATP per electron when acetate is the donor (E°’ of NO_3−_/NO_2−_ = +0.43 V *vs.* E°’ of Fe(III) (oxyhydr)oxides/Fe(II) ∼ −0.2 V *vs.* Standard Hydrogen Electrode (SHE)) [1, 2, 3]. As the redox potential of soils and sediments can vary widely [4, 5, 6], adjusting electron transfer chains to use acceptors with more favorable potentials allows anaerobes to maximize growth in response to environmental conditions [3, 7, 8, 9].

The respiration of Fe(III) and Mn(IV) poses unique challenges. These elements exist as insoluble (oxyhydr)oxides near neutral pH, requiring diversion of electrons from inner membrane respiratory chains to electron-accepting surfaces outside the cell [10, 11]. Additional complexity arises from the number of metal oxide polymorphs that exist in nature, with nearly 30 Mn oxides and 15 Fe oxides described, each with their own characteristic redox potential [12, 13, 14, 15]. While all of these could appear to the cell as similar extracellular electron sinks, the higher redox potential of Mn(IV) compared to Fe(III) oxides (E°’∼ +0.5 to +0.3 V for Mn(IV) *vs.* +0.1 to −0.3 V for Fe(III) *vs.* SHE) predicts that bacteria should be able to recognize and prefer specific metal forms. Sequential reduction of Mn(IV) before Fe(III) was observed in sediments as early as 1966 [5] and in pure cultures of *Geobacter metallireducens* in 1988 [16], suggesting that biological mechanisms exist to differentiate between higher *vs.* lower potential materials outside the cell.

*Geobacter spp.* can reduce multiple oxidized metals [17, 18, 19, 20], directly transfer electrons to methanogens [21], and utilize electrode surfaces as electron acceptors [22]. The complex array of electron transfer proteins in *Geobacter spp.* may explain this flexibility, with multiple *c*-type cytochromes and extracellular appendages identified that facilitate reduction of extracellular compounds. In *G. sulfurreducens*, at least five triheme cytochromes are linked to periplasmic electron transfer [23, 24], five multi-protein cytochrome complexes aid electron transfer through the outer membrane [25], and both multiheme cytochrome nanowires and extracellular pili extend beyond the cell [26, 27, 28]. Some outer membrane cytochromes are necessary for reduction of specific oxyanions such as SeO_32−_ [29], or use of Fe(III) *vs.* electrode surfaces [25, 30], but none explain how *Geobacter* might adapt its energy generation strategy to changes in redox potential.

The putative oxidoreductases ImcH and CbcL provide a possible mechanism for potential-dependent electron transfer [31, 32]. *G. sulfurreducens* requires the cytoplasmic membrane-localized seven-heme *c*-type cytochrome ImcH to respire extracellular acceptors above redox potentials of −0.1 V *vs.* SHE, and requires CbcL, a fusion of a diheme *b*-type cytochrome and a nine-heme *c*-type cytochrome, to use electron acceptors below −0.1 V *vs.* SHE. As *imcH* and *cbcL* are constitutively expressed [25, 32], the requirement for each appears to be controlled by ambient redox potential, somehow allowing cells to switch from ImcH-to CbcL-dependent electron transfer as conditions change [31, 32].

Multiple lines of evidence suggest ImcH and CbcL are not the only *G. sulfurreducens* oxidoreductases capable of routing electrons into the periplasm. The redox potentials of subsurface environments and microbial fuel cell anodes where *Geobacter spp.* typically dominate can be as low as −0.3 V *vs.* SHE, below the range where ImcH or CbcL are essential [33, 34]. Incubations of Δ*cbcL* with low-potential Fe(III) oxides such as goethite still produces Fe(II) [35], and Δ*cbcL* attached to electrodes still shows electron transfer below −0.2 V *vs.* SHE [32]. In addition, *Geobacter* genomes contain many uncharacterized gene clusters encoding a quinone oxidase-like *b*-type diheme cytochrome adjacent to a periplasmic multiheme *c*-type cytochrome, reminiscent of the two domains fused together in CbcL, and expression of some of these genes can be detected under metal-reducing conditions [36].

In this report, we identify CbcBA, a *bc*-type quinone oxidoreductase necessary for respiration near the thermodynamic limit of acetate oxidation. CbcBA is essential for extracellular metal and electrode reduction below −0.21 V *vs.* SHE, and is found within nearly every sequenced *Geobacter* genome [37]. We also provide evidence that use of CbcBA leads to lower growth yields, and may primarily act as a non-energy-conserving route for electron disposal. Unique from *imcH* and *cbcL*, *cbcBA* requires a σ^54^-dependent transcriptional activator for expression, and *cbcBA* is one of the most highly expressed genes during reduction of low potential Fe(III). Together, these cytochromes enable a branched electron transfer pathway that can operate at different redox potentials, allowing ImcH-dependent respiration when potential energy is plentiful, CbcL-dependent growth as energy becomes limiting, and use of CbcBA near the threshold able to support microbial life.

## Materials and Methods

### Bacterial strains and culture conditions

All strains and plasmids used in this study are listed in Table 1. *G. sulfurreducens* strains and mutants were grown in a minimal medium (referred to as NB) containing 0.38 g.L^−1^ KCl, 0.2 g.L^−1^ NH_4_Cl, 0.069 g.L^−1^ NaH_2_PO_4_.H_2_O, 0.04 g.L^−1^ CaCl_2_.2H_2_O, 0.2 g.L^−1^ MgSO_4_.7H_2_O, 10 mL of trace mineral mix, and buffered with 2 g.L^−1^ of NaHCO_3_ purged with N_2_:CO_2_ (80:20) atmosphere, incubated at 30 °C. Trace mineral mix was composed of 1.5 g.L^−1^ nitrilotriacetic acid as a chelator for growth, except when grown with Fe(III)-oxides, in which case minerals were prepared in 12.5 mL.L^−1^ of 7.7 M HCl to a final concentration of 0.1 M HCl, 0.1 g.L^−1^ MnCl_2_.4H_2_O, 0.5 g.L^−1^ FeSO_4_.7H_2_O, 0.17 g.L^−1^ CoCl_2_.6H_2_O, 0.10 g.L^−1^ ZnCl_2_, 0.03 g.L^−1^ CuSO_4_.5H_2_O, 0.005 g.L^−1^ AlK(SO_4_)_2_.12H_2_O, 0.005 g.L^−1^ H_3_BO_3_, 0.09 g.L^−1^ Na_2_MoO_4_, 0.05 g.L^−1^ NiCl_2_, 0.02 g.L^−1^ Na_2_WO_4_.2H_2_O, 0.10 g.L^−1^ Na_2_SeO_4_. Routine growth was performed in acetate-fumarate NB medium (NBFA) containing 20 mM acetate as the carbon source and electron donor and 40 mM fumarate as the electron acceptor. For solid medium, 1.5% agar was added to acetate-fumarate medium for growth on plates in an anaerobic workstation (Microbiology International, Maryland) under N_2_: CO_2_: H_2_ (75:20:5) atmosphere maintained at 30 °C. Every experiment was initiated by streaking fresh strains of *G. sulfurreducens* from −80 °C culture stocks. 200 µg.mL^−1^ kanamycin was used for *G. sulfurreducens*, 100 µg.mL^−1^ ampicillin and 50 µg.mL^−1^ kanamycin for *Escherichia coli* as indicated.

**Table 1:** List of strains and plasmids used in this study.

| Strains or Plasmids | Description or relevant genotype | Reference or source |
| --- | --- | --- |
| ***G. sulfurreducens* strains** |  |  |
|  | Wildtype *G. sulfurreducens* (WT) | Lab culture collection |
| DB789 | $\Delta imcH$ | [38] |
| DB868 | $\Delta cbcL$ | [32] |
| DB790 | $\Delta imcH \Delta cbcL$ | This study |
| DB1674 | $\Delta bccR$ | This study |
| DB1717 | $\Delta cbcBA$ | This study |
| DB1718 | $\Delta imcH \Delta cbcBA$ | This study |
| DB1719 | $\Delta cbcL \Delta cbcBA$ | This study |
| DB1720 | $\Delta imcH \Delta cbcL \Delta cbcBA$ | This study |
| DB1721 | $\Delta cbcBA$ Tn7::p0597 *cbcBA*-Kan | This study |
| DB1725 | $\Delta cbcBA$ Tn7::p0597 *cbcB*-Kan | This study |
| DB1729 | $\Delta cbcBA$ Tn7::p0597 *cbcA*-Kan | This study |
| ***E. coli* strains** |  |  |
| UQ950 | Cloning strain of *E. coli* |  |
| S17-1 | Conjugation donor strain |  |
| MFDpir | Conjugation donor strain |  |
| DB2068 | S17-1 strain containing pk18mobsacBDGSU0593-94 for *cbcBA* deletion | This study |
| DB1658 | S17-1 strain containing pk18mobsacBDGSU0598 for *bccR* deletion | This study |
| DB1325 | MFDpir conjugation donor strain containing helper plasmid pmobile-CRISPRi | [85] |
| DB2075 | UQ950 strain containing pGeo2::p0597 *cbcBA* | This study |
| DB2076 | UQ950 strain containing pGeo2::p0597 *cbcB* | This study |
| DB2077 | UQ950 strain containing pGeo2::p0597 *cbcA* | This study |
| DB2079 | UQ950 strain containing pTn7c147::p0597 *cbcBA* | This study |
| DB2081 | UQ950 strain containing pTn7c147::p0597 *cbcB* | This study |
| DB2083 | UQ950 strain containing pTn7c147::p0597*cbcA* | This study |
| **Plasmids** |  |  |
| pk18mobsacB |  |  |
| prk2Geo2 |  | [38] |
| pTn7C147 |  | [39] |
| pmobile-CRISPRi |  | [85] |
| pDGSU0593-94 | Flanking regions of GSU0593-GSU0594 in pk18mobsacB | This study |
| pDGSU0598 | pk18mobsacB deletion vector containing flanking regions of GSU0598. | This study |
| prk2Geo2::0597 *pcbcBA* | Complementation vector of GSU0593-0594 with its predicted native promoter | This study |
| prk2Geo2::0597 *pcbcB* | Complementation vector of GSU0593 with its predicted native promoter | This study |
| prk2Geo2::0597 *pcbcA* | Complementation vector of GSU0594 with its predicted native promoter | This study |
| pTn7C147::p0597 *cbcBA*-Kan | Complementation vector subcloned from pGeo2 expressing *cbcBA* under the control of native promoter. | This study |
| pTn7C147::p0597 *cbcB*-Kan | Complementation vector subcloned from pGeo2 expressing *cbcB* under the control of native promoter. | This study |
| pTn7C147::p0597 *cbcA*-Kan | Complementation vector subcloned from pGeo2 expressing *cbcA* under the control of native promoter. | This study |

### Strain construction and complementation

Deletion constructs were designed based on a strategy previously described [38]. Briefly, ∼1 kb upstream and downstream region of *cbcBA* (GSU0593-0594), and *bccR* (GSU0598) were amplified using primers listed in Table 2. Amplified upstream and downstream DNA fragments were fused using overlap extension PCR. Amplified fused DNA fragments were digested with restriction enzymes listed in Table 2, and ligated into digested and gel purified pk18mobsacB. The ligation product was transformed into UQ950 chemically competent cells. The resulting plasmid was sequence verified before transformation into S17-1 conjugation donor cells. Overnight grown S17-1 donor strain containing the plasmid was conjugated with *G. sulfurreducens* acceptor strain inside an anaerobic chamber on a sterile filter paper placed on an NBFA agar plate. After ∼4 h, cells scraped from filters were streaked on NBFA agar plates containing kanamycin. The positive integrants were streaked on NBFA + 10% sucrose plates to select for the wildtype or deletion genotype. Colonies from NBFA + 10% sucrose plates were patched on NBFA and NBFA + 200 µg.mL^−1^ to identify antibiotic sensitive, markerless deletion strains. The strains were verified by PCR for the gene deletion and final strains checked for off-site mutations via Illumina re-sequencing.

**Table 2:**
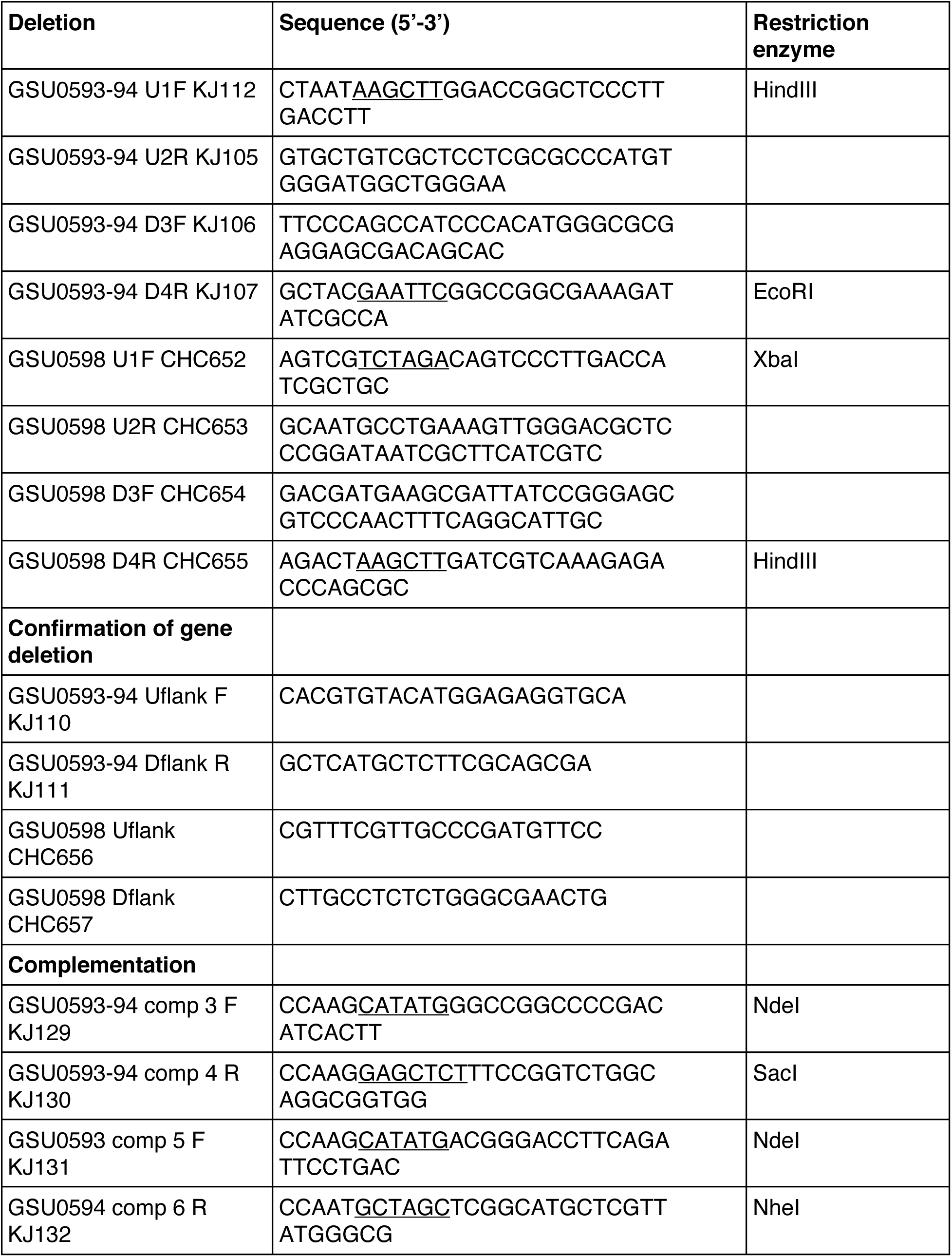
Primers used in this study.

Complementation was performed using the method described in Hallberg *et al.* [39]. Complement strains were constructed by first cloning *cbcBA* (GSU0593-94), *cbcB* (GSU0593), or *cbcA* (GSU0594) gene into the pRK2Geo2 vector. The *cbcBA* cluster with native ribosomal binding sites was cloned under the control of its native promoter (GSU0597). The resulting vectors were sequence verified, then subcloned into pTn7c147 between the n7L and n7R regions. Newly subcloned pTn7 vectors were transformed in MFDpir chemically competent cells [40]. Any DNA between n7L and n7R regions is integrated downstream of the *glmS* (GSU0270) site, surrounded by strong terminators [41]. A helper plasmid pJMP1039 (a derivative of pTNS3) expressing recombinase TnsABCD in MFDpir cells was utilized to recognize n7L and n7R regions in pTn7 vectors [41], and integrate DNA onto *G. sulfurreducens* chromosome downstream of *glmS.* A triparental mating strategy was used to create complement strains.

Integrating genes onto the genome minimizes growth-rate and biofilm defects encountered when using most plasmids in *G. sulfurreducens*.

### Cyclic voltammetry

Three-electrode bioreactors contained 3 cm^2^ 1500-grit polished polycrystalline graphite working electrodes (POCO AXF-5Q, TriGemini LLC, Illinois), platinum wire counter electrodes, Ag/AgCl reference electrodes [42, 43], and were autoclaved at 121 °C for 20 min. Anoxic conditions were maintained by constantly flushing reactors with anoxic humidified N_2_: CO_2_ (80:20) gas. Acetate (40 mM) served as the electron donor and carbon source, and poised electrodes (+0.24 V *vs.* SHE) served as the electron acceptor. Acetate-fumarate grown cells (acceptor limited, OD_600_ ≃ 0.5) were inoculated at 25% v/v inoculation into 30 °C stirred reactors. A 16-channel potentiostat (Biologic Science Instruments, France) constantly recorded anodic current over time. Cyclic voltammetry was applied by forward scanning electrode potential from −0.55 V *vs.* SHE to +0.24 V *vs.* SHE, and reverse scanned back to −0.55 V *vs.* SHE at 1 mV/s for two scans [42].

### Growth with Fe(III) citrate

Minimal medium containing 20 mM acetate and 55 mM Fe(III) citrate was used in anaerobic Balch tubes, or in bioreactors when redox potential was measured over time [35, 43]. Media were autoclaved at 121 °C and immediately removed to cool at room temperature in the dark. Anaerobic tubes containing Fe(III) citrate medium were inoculated at 1:100 v/v from stationary phase cultures (OD_600_ ≃ 0.5) grown in NBFA. 0.1 mL of sample was taken at regular intervals and dissolved in 0.9 mL of 0.5 N HCl. Fe(II) concentrations were measured using a ferrozine assay [44].

### Redox potential measurement

For monitoring redox potential, bioreactors were used in Open Circuit Potential (OCP) mode. Short (1cm) electrochemically cleaned platinum wires were used as sensing electrodes with a Ag/AgCl reference (+0.21 V *vs.* SHE). Platinum was cleaned in 0.5 M H_2_SO_4_ by holding the working electrode at +2.24 V *vs.* SHE, cycling electrode potential between +0.01 V and +1.34 V for 20 cycles and stopping at +1.34 V *vs.* SHE.

### Fe(III) oxide reduction

Medium containing 20 mM acetate and either ∼50 mM akaganeite or ∼30 mM hydrous ferric oxide was supplemented with 0.69 g.L^−1^ NaH_2_PO_4_.H_2_O (to prevent formation of crystalline Fe(III) (oxyhydr)oxide while autoclaving) [14]. Fresh akaganeite was synthesized by slowly adding 25% NaOH dropwise over the course of 1 h into a stirring solution of 0.4 M FeCl_3_ to pH 7. The suspension was aged for at least one hour at pH 7, then washed with DI H_2_O via centrifugation. 1 mL of freshly synthesized akaganeite (β-FeOOH) (∼0.5 M) was added to 9 mL medium with 20 mM acetate as the carbon source before autoclaving [14, 35]. Hydrous ferric oxide was synthesized first as schwertmannite (Fe_8_O_8_(OH)_6_(SO_4_).nH_2_O) by adding 5.5 mL of 30% hydrogen peroxide to a solution of 10 g.L^−1^ FeSO_4_, then stirred overnight to stabilize. Schwertmannite solids were washed with DI H_2_O thrice by centrifugation. The resulting mineral was added to medium with 20 mM acetate as the carbon source before autoclaving. Autoclaving at neutral pH transforms the schwertmannite into ferrihydrite with an amorphous XRD-signature [14, 35, 39]. Iron oxide medium was inoculated with 1:100 v/v of cells (OD_600_ ≃ 0.5) grown in NBFA medium. Samples (0.1 mL) were dissolved in 0.9 mL 0.5 N HCl, and stored in the dark before measurement via ferrozine assay.

### Transcriptomic analysis using RNA-seq

Total RNA was extracted from *G. sulfurreducens* fumarate-grown cultures in exponential phase. For cells grown with Fe(III) citrate, RNA was extracted from cultures at exponential growth phase when ∼30% or ∼70% of Fe(III) citrate was reduced. Cells were collected using vacuum filtration to minimize inhibition from Fe(III)/Fe(II) in the medium. Electrode biofilms were scraped from electrodes immediately after disconnecting them from the potentiostat. All cell pellets were washed in RNAprotect reagent (Qiagen) and stored at −80 °C before extraction using RNeasy with on column DNase treatment (Qiagen). Ribosomal RNA was depleted using Ribozero (Illumina) before sequencing on the Illumina Hiseq 2500 platform in 125-bp pair-ended mode. Residual ribosomal RNA sequences were removed using Bowtie2 [45] before analysis. Duplicate rRNA-depleted biological samples were analyzed for each strain and condition using Rockhopper [46], with our re-sequenced *G. sulfurreducens* genome as reference [38]. Expression was normalized by reads mapped by the upper quartile of gene expression values, and full RNA-seq data are in Supplementary table 1.

### CFU and yield measurements

Growth of *G. sulfurreducens* strains was measured by counting colony-forming units (CFUs). A drop plate method adapted from Herigstad *et al*. [47], was used to count cells on NBFA agar medium. Briefly, 100 μL of samples were serially diluted 1:10 in liquid medium, and 10 µL of each dilution was plated on NBFA agar plates inside an anaerobic chamber (Coy laboratory products, Michigan) with an N_2_: CO_2_: H_2_ (75:20:5) atmosphere. Total Fe(III) reduced was measured using a ferrozine assay, so cellular yield could be calculated as CFU per mM Fe(III) reduced as cells were actively growing.

## Results

### The *cbcBA* gene cluster encodes a *b*-and *c*-type cytochrome expressed late in Fe(III) reduction

The *G. sulfurreducens* genome contains at least six putative inner membrane quinone oxidoreductase gene clusters. Five encode both *b*- and *c*-type cytochrome domains: Cbc1 (GSU0274, *cbcL*), Cbc3 (GSU1648-GSU1650, *cbcVWX*), Cbc4 (GSU0068-GSU0070, *cbcSTU*), Cbc5 (GSU0590-GSU0594, *cbcEDCBA*), Cbc6 (GSU2930-GSU2935, *cbcMNOPQR*) [48], and one contains only a multiheme *c*-type cytochrome (GSU3259, *imcH*) [31]. The *b-* and *c-*type cytochrome CbcL (Cbc1) is essential for growth below redox potentials of about −0.1 V *vs.* SHE [32], while the *c*-type cytochrome ImcH is essential for respiration as redox potential rises above this point [31]. Among these *b*- and *c*-type cytochrome gene clusters, Cbc5 is the most conserved cytochrome-containing gene cluster among *Geobacter* species [37].

Bioinformatic [49, 50, 51] and transcriptomic analyses [25, 52] place *cbcBA* in an operon with a σ^54^-dependent promoter upstream of GSU0597 and a transcriptional terminator downstream of *cbcB* (Figure 1A). This operon encodes two hypothetical proteins (GSU0597 and GSU3489), a RpoN-dependent response regulator (GSU0596), a quinone oxidoreductase-like di-heme *b*-type cytochrome (CbcB) [53], and a seven-heme *c*-type cytochrome (CbcA) (Figure 1C). An inner membrane localization of CbcBA is predicted by PSORT [54], with CbcB integrated into the inner membrane and CbcA exposed in the periplasm anchored by a C-terminal transmembrane domain. Cell fractionation studies also report a cytoplasmic membrane association of CbcA [55], implying that CbcBA is located at the inner membrane.

**Figure 1.**
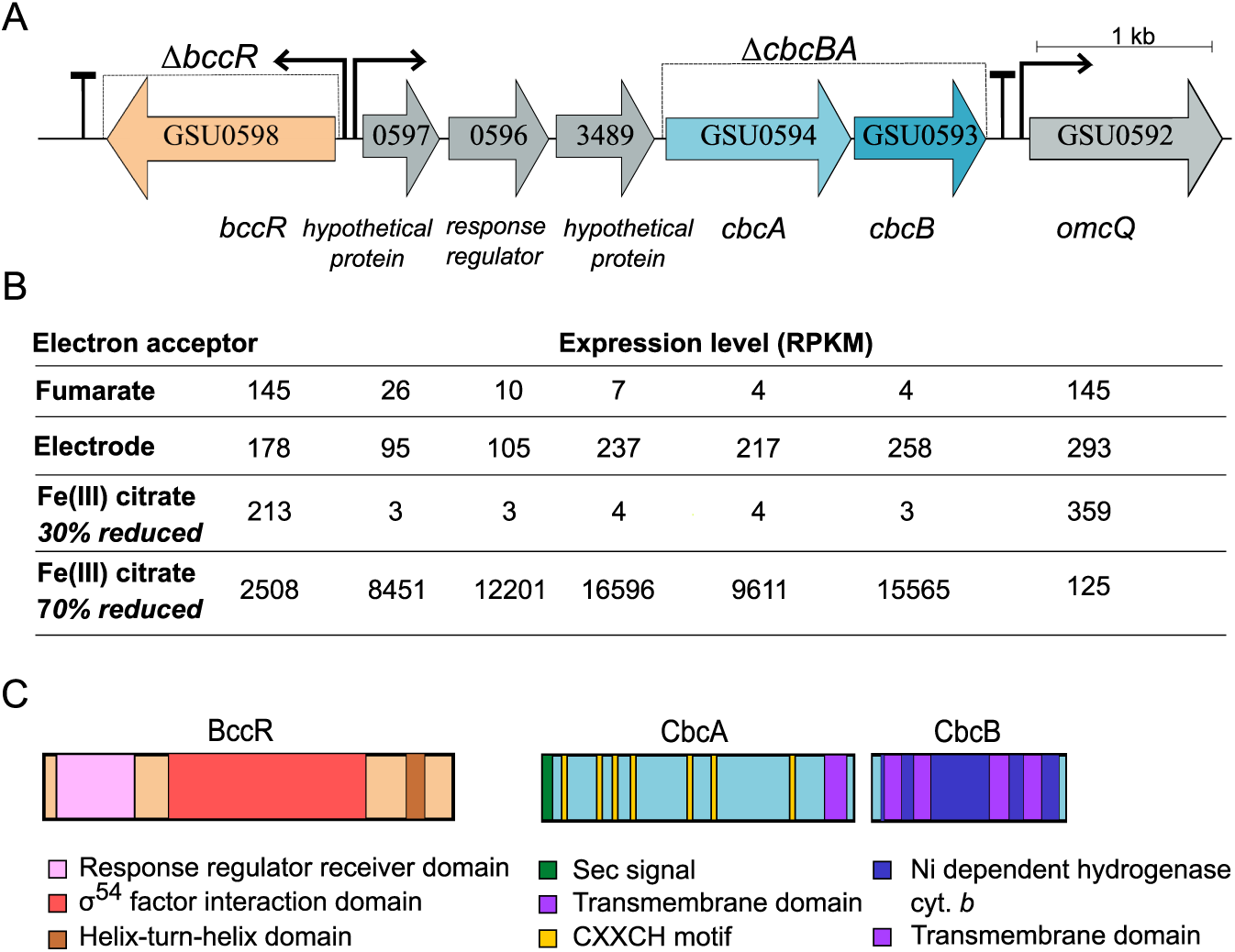
Genetic organization, expression, and domain structure of the *cbcBA* region. A. *cbcBA* is of a five-gene operon divergently transcribed from a σ54-dependent transcriptional regulator (*bccR*). B. Expression levels, as reads per kilobase mapped (RPKM), when growing exponentially with different electron acceptors: fumarate, electrodes, or Fe(III) citrate (30% *vs*. 70% Fe(III) reduced). The expression values correspond to the gene labels in panel A. See Supplementary Table 1 for full RNA-seq data. C. Predicted features and domain structure of BccR, CbcB, and CbcA.

Divergently transcribed from this operon is GSU0598, a putative σ^54^-dependent transcriptional regulator, which we have named *bccR* (for *<u>bc</u>*-type <u>c</u>ytochrome regulator) (Figure 1A). BccR belongs to the RpoN-dependent family of regulators that bind −12 /−24 elements [56]. BccR contains a response receiver domain, a σ^54^ factor interaction domain, and a C-terminal helix-turn-helix domain [57] (Figure 1C).

The *cbcBA* operon (GSU0597-GSU0593) had near zero expression when fumarate was the electron acceptor, but low expression was detected in electrode-grown biofilms [25] (Figure 1B). When growing with Fe(III) citrate as the electron acceptor, expression of the *cbcBA* operon remained low during the first 20 h of growth (Figure 1B), or as the first ∼30% Fe(III) was reduced (Figure 2A). However, *cbcBA* was dramatically upregulated after 30 h of growth (Figure 1B), as ∼70% of Fe(III) became reduced (Figure 2A). The level of *cbcBA* expression (>12 000 RPKM) was higher than 99% of *G. sulfurreducens* genes at this stage (SI figure 1).

**Figure 2.**
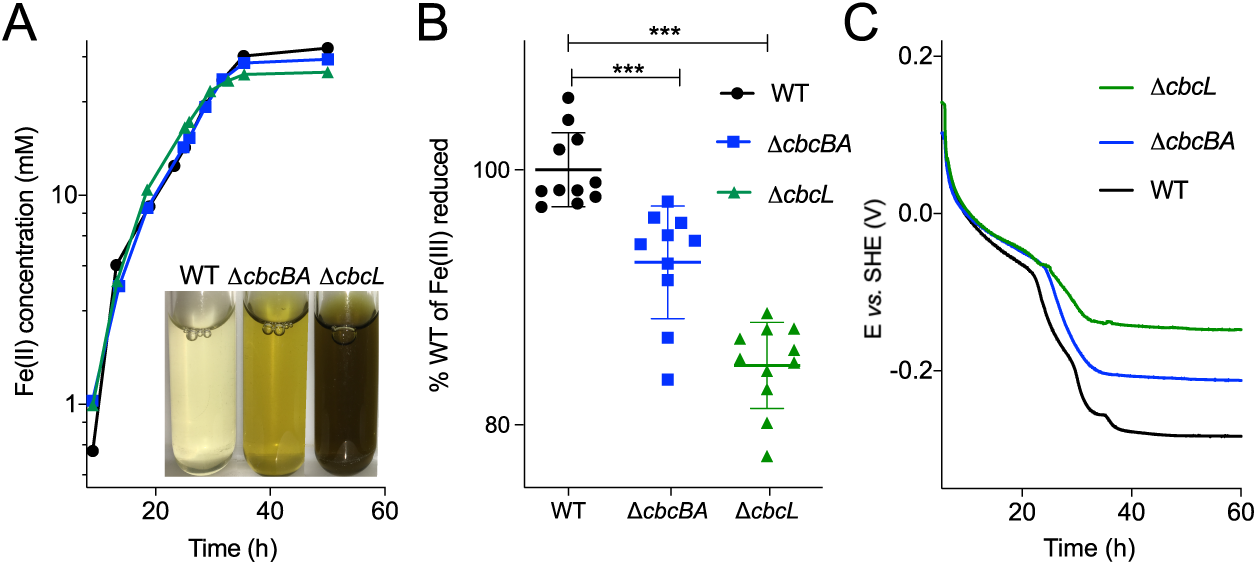
Mutants lacking *cbcBA* or *cbcL* cannot reduce all available Fe(III) citrate, and *cbcBA* is required for reduction below −0.21 V *vs*. SHE. A. Fe(III) citrate reduction over time. Inset image shows the difference in endpoint Fe(III) citrate reduction by different strains. B. Δ*cbcBA* reduces ∼93% of Fe(III) citrate compared to WT, whereas Δ*cbcL* reduces ∼85% of Fe(III) citrate compared to WT. C. Redox potential recorded over time in the same medium, as cells reduce Fe(III) citrate. The Δ*cbcL* mutant fails to lower redox potential below −0.15 V *vs*. SHE whereas the Δ*cbcBA* mutant fails to lower redox potential below −0.21 V vs. SHE. All experiments were conducted in triplicate, and representative curves are shown in A and C (N ≥ 5). B shows end point values from individual experiments averaged with standard deviation reported as error bars (n ≥ 10). Two-tailed t-test was performed to calculate p-values.

### CbcBA is essential for complete reduction of Fe(III) citrate

To determine if CbcBA was involved in extracellular electron transfer, a markerless deletion of GSU0593-94 (Δ*cbcBA*) was created. The Δ*cbcBA* mutant did not show any defect with fumarate as the electron acceptor (SI figure 2). However, the extent of Fe(III) reduction by Δ*cbcBA* was lower. Mutants lacking *cbcBA* never reduced the final 8-10% of Fe(III) citrate (Figure 2A), regardless of the amount of electron donor provided or length of incubation.

The putative quinone oxidoreductase ImcH is essential for reduction of high potential electron acceptors such as freshly prepared Fe(III) citrate [31], while the *bc*-cytochrome CbcL becomes essential as Fe(III) is reduced and redox potential drops [32, 35]. As the type of Fe(III) reduction defect observed for Δ*cbcBA* was similar to Δ*cbcL*, mutants lacking *cbcL* and *cbcBA* were directly compared. The Δ*cbcBA* strain ceased reduction of Fe(III) after 92.7 ± 1.4% (n=10) of Fe(III) citrate was reduced, whereas Δ*cbcL* only reduced 84.6 ± 1.0% (n=11) of Fe(III) citrate (Figure 2A, 2B). This suggested that CbcBA became necessary in the final stages of Fe(III) reduction.

### CbcBA is required for reduction of Fe(III) citrate below −0.21 V *vs.* SHE

Because the >3000-fold up-regulation of *cbcBA* occurred after more than half of Fe(III) citrate was reduced (SI figure 1), induction of *cbcBA* did not appear to be due to the presence of Fe(III) *per se*. To more accurately determine the energy available during each stage of Fe(III) reduction, we measured redox potential continuously during growth with a potentiostat [35, 58]. Redox potential titrations and voltammetry determined the midpoint potential of the Fe(II) citrate/Fe(III) citrate half-reaction in our medium to be −0.043 V *vs.* SHE (SI figure 3). This is lower than values calculated in literature, likely due to high levels of chelating carboxylic acids in commercial Fe(III) citrate combined with electron donors, creating bi- or tri-dentate complexes with lower redox potential than the 1:1 ratios assumed in standard calculations [59, 60, 61].

When wildtype (WT) cells were inoculated into freshly prepared Fe(III) citrate (>99% oxidized), redox potential dropped rapidly from +0.15 V, and stabilized days later at −0.27 V *vs.* SHE when nearly 100% of Fe(III) was reduced. Considering the formal redox potential of CO_2_/acetate is −0.28 V, cells utilized nearly all the free energy available. In contrast, Δ*cbcL* ceased Fe(III) reduction near −0.15 V *vs.* SHE (equivalent to 38 mM Fe(III) reduced) [35]. Under the same conditions, Δ*cbcBA* stabilized at −0.21 V *vs.* SHE (equivalent to ∼46 mM Fe(III) reduced). Each mutant produced these same endpoint potentials independent of inoculation size or incubation time (SI figure 4), or when the concentration of Fe(III) citrate was increased to 80 mM [35].

### Complementation of Δ*cbcBA* requires both *cbcB* and *cbcA*

To test if *cbcB* or *cbcA* alone were responsible for this inability to reduce Fe(III) below −0.21 V *vs.* SHE, single genes were integrated into the chromosome under control of the *cbcBA* operon’s promoter [39]. When Δ*cbcBA*::*cbcB* or Δ*cbcBA*::*cbcA* strains were grown with Fe(III) citrate, reduction still ceased at the same extent and redox potential (Figure 3A).

**Figure 3.**
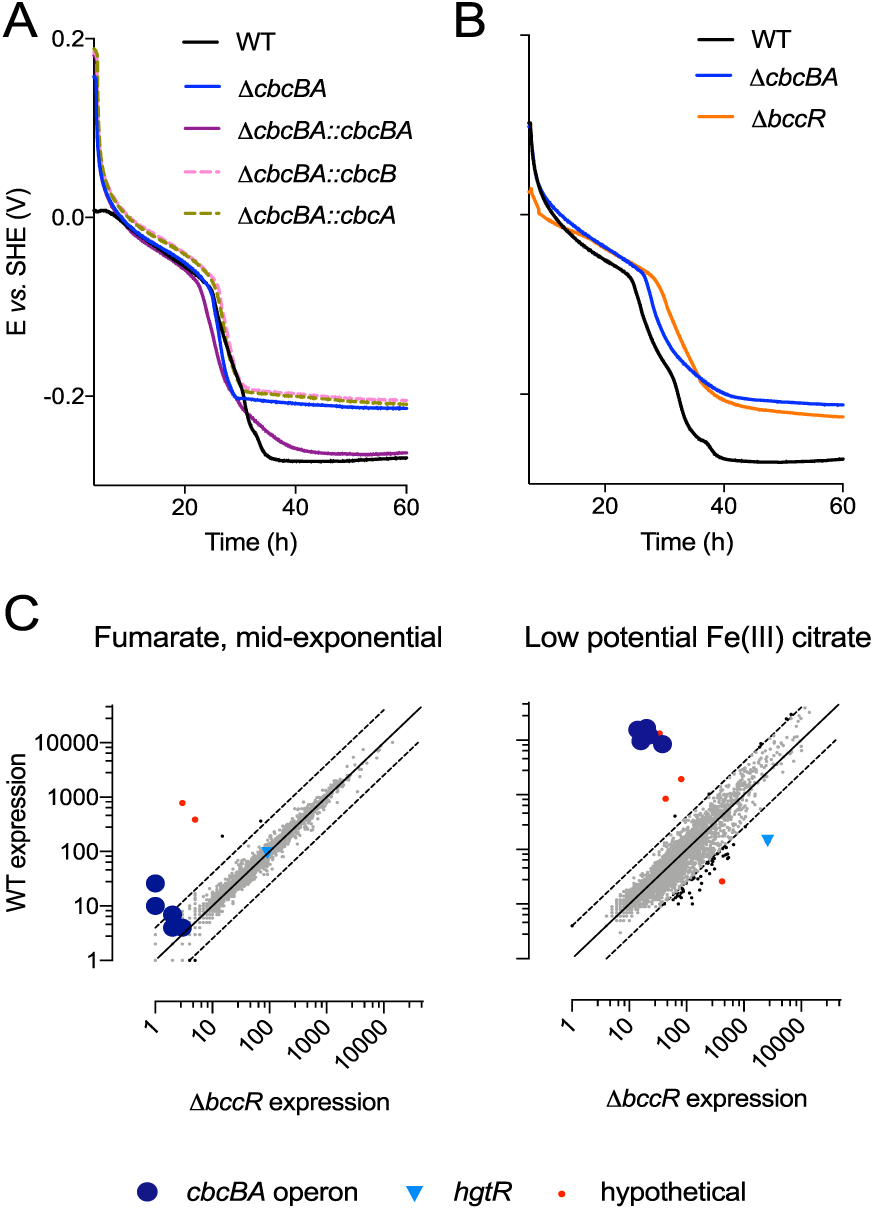
Complementation of Δ*cbcBA* requires expression of both *cbcB* and *cbcA*, and *bccR* essential for induction of *cbcBA*. A. Complementation with both *cbcB* and *cbcA* are required to fully restore Fe(III) reduction in Δ*cbcBA*. B. Deletion of *bccR*, a σ^54^-dependent transcriptional response regulator upstream of the *CbcBA* operon and comparison with Δ*cbcBA*. C. Transcriptomic analysis of WT *G. sulfurreducens vs*. Δ*bccR* grown with fumarate or Fe(III) citrate. Points above the 1:1 line indicate reduced expression due to deletion of *bccR*, points below the line had increased expression when *bccR* was deleted.

However, when both *cbcBA* genes were integrated and expressed in the Δ*cbcBA* strain, the extent of Fe(III) reduction was restored to WT levels (Figure 3A). Based on these results, all subsequent experiments were conducted with mutants lacking both genes. **BccR is necessary for expression of *cbcBA.*** A response regulator is divergently transcribed upstream of *cbcBA bc*-type cytochrome operons in all examined *Geobacter* genomes [25, 62]. When *bccR* (GSU0598) was deleted, Δ*bccR* ceased reduction of Fe(III) at −0.21 V *vs.* SHE, the same potential as Δ*cbcBA* (Figure 3B). RNAseq revealed that expression of *cbcBA* was no longer upregulated in Δ*bccR* during Fe(III) citrate reduction (Figure 3C) consistent with BccR being an activator of the *cbcBA* operon. Deletion of *bccR* did not affect other putative quinone oxidoreductases, in particular *imcH* or *cbcL*, which were constitutively expressed at much more moderate (∼500 RPKM) levels.

While the largest effect of *bccR* deletion was downregulation of *cbcBA* operon (Figure 3C), Δ*bccR* showed upregulation of *hgtR* (GSU3364) when Fe(III) was the electron acceptor. HgtR is a RpoN-dependent repressor involved in downregulating acetate oxidation when hydrogen is the electron donor [56, 63]. The increase in *hgtR* expression by more than 1 000x in Δ*bccR* implies a possible role for HgtR in down-regulating the TCA cycle during reduction of Fe(III) as acetate oxidation becomes thermodynamically limited.

### Double mutants show that *imcH*, *cbcL*, and *cbcBA* are required within different redox potential windows

If one inner membrane cytochrome is needed in order to lower redox potential enough to activate the next, then double and triple markerless deletion mutant strains should still show the phenotype of their dominant missing pathway. All single, double, and triple mutant strains lacking *imcH* failed to initiate Fe(III) citrate reduction when inoculated into fresh >+0.1 V *vs.* SHE medium, and did not lower the redox potential more than 20 mV over the following 60 h (Figure 4). The dominance of Δ*imcH* in all backgrounds corroborates data showing ImcH to be essential for electron transfer in fresh Fe(III) citrate, Mn(IV) oxide, and electrodes at redox potentials above 0 V [31, 35], and showed that the presence or absence of *cbcBA* did not alter this behavior.

**Figure 4.**
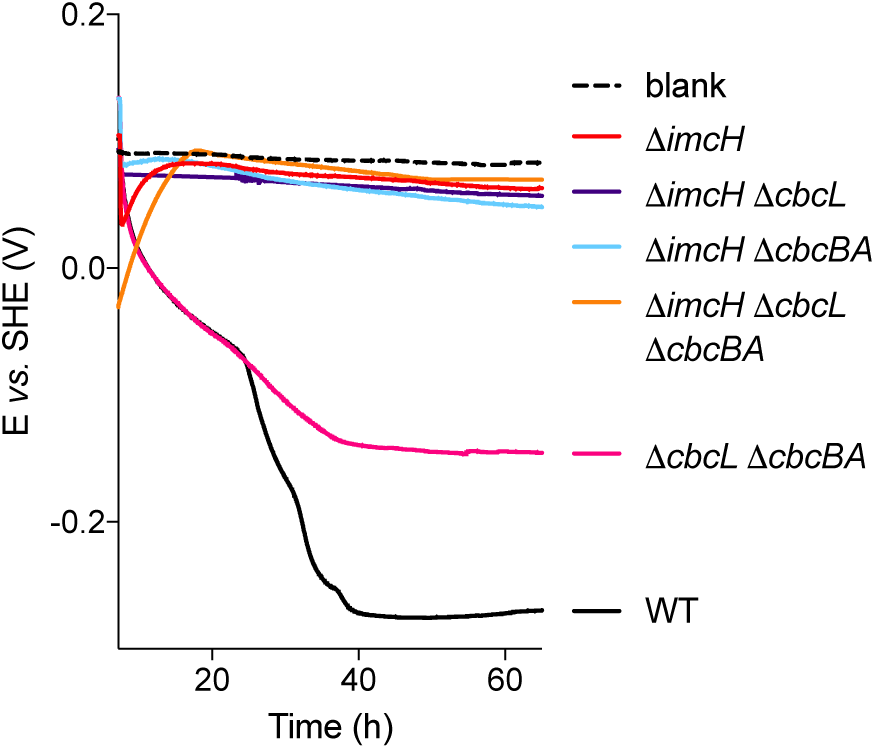
Deletion of *imcH* prevents reduction of high potential Fe(III) citrate in all mutant backgrounds, while deletion of *cbcL* always prevents reduction beyond −0.15 V *vs*. SHE. Mutants lacking *imcH* (Δ*imcH*, Δ*imcH* Δ*cbcL*, Δ*imcH* Δ*cbcBA*, Δ*imcH* Δ*cbcL* Δ*cbcBA*) fail to reduce fresh Fe(III) citrate and cannot lower redox potential. The double mutant lacking *cbcL* and *cbcBA* (Δ*cbcL* Δ*cbcBA*) fails to lower redox potential below −0.15 V *vs*. SHE, similar to the Δ*cbcL* single mutant. Representative curves from experiments conducted in triplicates are shown here.

Like the single Δ*cbcL* mutant, the Δ*cbcL* Δ*cbcBA* double mutant containing *imcH* initially reduced Fe(III), then ceased reduction at −0.15 V *vs.* SHE. This provides additional evidence that ImcH can function down to a redox potential of −0.15 V, and that only CbcL can lower redox potential beyond this point, regardless of whether CbcBA is present (Figure 4). The phenotype of Δ*cbcBA* (Figure 2, 3) similarly implies that CbcL is essential until −0.21 V *vs.* SHE, at which point CbcBA is required for electron transfer (Figure 3).

### Cyclic voltammetry detects a CbcBA-dependent electron transfer process with a midpoint potential of −0.24 V *vs.* SHE

All evidence up to this point that *cbcBA* was required at specific redox potentials was derived from soluble Fe(III) incubations, which could be non-physiological compared to environments where *G. sulfurreducens* uses a partner in syntrophy, or a solid electrode as the electron acceptor. To examine electron transfer in the absence of Fe(III), we grew *G. sulfurreducens* on graphite electrodes, and subjected the biofilms to cyclic voltammetry. During cyclic voltammetry, redox potential can be brought to a value too low to support acetate oxidation (−0.4 V *vs.* SHE) to obtain a baseline. When electrode potential is slowly increased, electron transfer from adherent cells is observed at a key onset potential as it becomes thermodynamically favorable, accelerates until a maximum electron transfer rate is reached, and follows the reverse trend as potential is decreased.

In theory, when a single event is rate-limiting in voltammetry, a Nernstian sigmoidal rise in current occurs over a ∼100 mV window, rising most steeply at the potential that most strongly affects the oxidation state of a key redox center. The potential-dependent responses of *G. sulfurreducens* cells during voltammetry are more complex than one-event models, and instead display at least three overlapping processes [32, 42, 64, 65, 66]. These three inflection points can be easily identified by displaying the first derivative of current increase as a function of applied potential.

Prior work described a change in voltammetry near −0.10 V *vs.* SHE when *cbcL* was deleted, which could be restored by *cbcL* complementation [32]. These experiments also detected a lower potential process independent of CbcL that increased with each subsequent voltammetry sweep. Impedance measurements by Yoho *et al.* [65] reported a similar low potential electron transfer process detectable within minutes of applying reducing electrode potentials. Based on our data, we hypothesized these unexplained features [32, 65] could be due to *cbcBA* activation during exposure to low potential electrodes.

To test this hypothesis, we first grew WT and Δ*cbcL* biofilms on electrodes as electron acceptors at +0.24 V *vs.* SHE, then subjected biofilms to voltammetry sweeps to reveal the low potential response below −0.2 V, and confirm loss of the middle −0.1 to −0.15 V process attributed to CbcL (Figure 5A). When *cbcBA* was deleted in the Δ*cbcL* background, the low potential electron transfer event disappeared, and all electron transfer below −0.15 V was eliminated. In the single Δ*cbcBA* mutant, only current below −0.2 V was eliminated, further linking *cbcBA* to activity in this low potential range (Figure 5A). By plotting the first derivative of voltammetry data, regions where changes in potential caused the steepest response(s) could be identified. According to these data, deletion of *cbcBA* eliminated an electron transfer process between −0.28 and −0.21 V, with a midpoint potential of −0.24 V *vs.* SHE.

**Figure 5.**
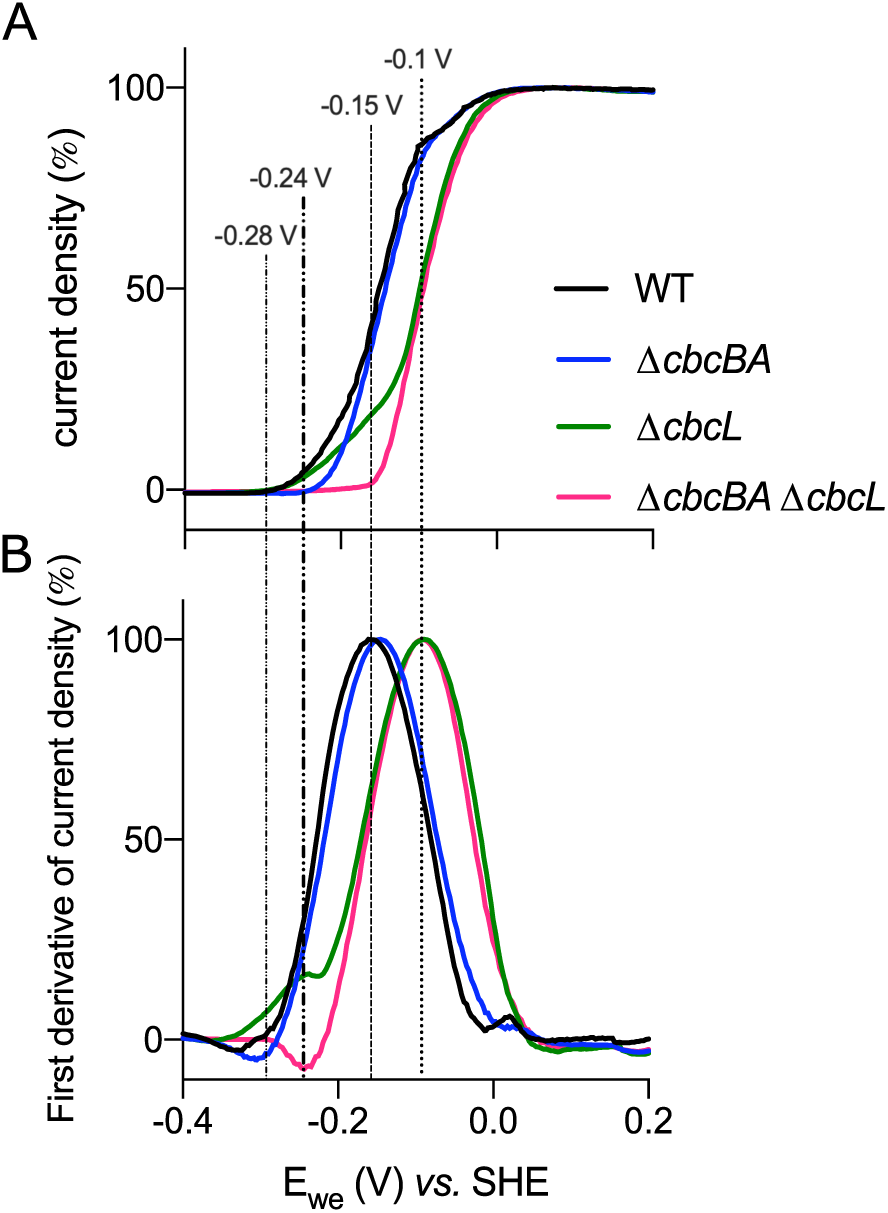
Activation of a CbcBA-dependent electron transfer pathway at redox potentials below −0.21 V vs. SHE. A. *G. sulfurreducens* mutants grown on poised electrodes (+0.24 V *vs.* SHE) to 300-400 μA.cm^−2^ subjected to cyclic voltammetry at 1 mV/s scan rate in the presence of acetate. The Δ*cbcL* mutant still showed a WT-like onset potential at −0.28 V *vs.* SHE, and retained electron transfer at potentials below −0.2 V. The Δ*cbcBA* mutant lost this low potential ability and shifted to a more positive onset potential. B. First derivative of cyclic voltammetry data of mutant strains revealed clear differences in the potential where maximal rate of reduction occurs. The Δ*cbcL* strain lacked the WT response at −0.15 V *vs.* SHE, corresponding to reduction defect of low potential electron acceptors. The Δ*cbcBA* Δ*cbcL* double mutant lacked another low potential response at −0.24 V *vs.* SHE. All experiments were conducted in duplicate, two scans were performed, and data from the reverse second scan was used for analysis.

### CbcBA is essential for complete reduction of different Fe(III) (oxyhydr)oxides

With the evidence that *G. sulfurreducens* not only required *cbcBA* for electron transfer to soluble metals, but also to electrode surfaces, we then asked if *cbcBA* was involved in reduction of insoluble Fe(III) (oxyhydr)oxide particles found in the environment [13]. While common forms such as ferrihydrite, akaganeite, goethite, and hematite all have the same chemical formula (FeOOH), these minerals differ greatly in calculated redox potentials [67]. For example, freshly synthesized hydrous ferric oxide possesses a relatively high redox potential (+0.1 to 0 V, depending on age and surface area) [15, 68], while more crystalline hematite can be as low as −0.2 to −0.3 V [69]. These differences could affect the relative importance of *cbcBA*, especially if a lower-potential form is available.

To compare insoluble Fe(III) minerals, two different forms representing progressively lower redox potential acceptors compared to Fe(III) citrate were synthesized. First, single mutants were incubated with a freshly precipitated hydrous ferric oxide, which has an estimated redox potential of ∼0 V *vs.* SHE. Consistent with this acceptor having a potential near where both ImcH and CbcL have both been shown to operate, Δ*imcH* initially reduced Fe(III) slowly, until Fe(II) accumulated to 1-2 mM, then accelerated to reduce nearly the same total Fe(III) as reduced by WT cells (Figure 6A). The mutant lacking *cbcL* reduced only 50% of Fe(III), and Δ*cbcBA* reduced 90% of total Fe(III) compared to WT (Figure 6A). This pattern was similar to Fe(III) citrate, but showed increased importance of both *cbcL* and *cbcBA*.

**Figure 6.**
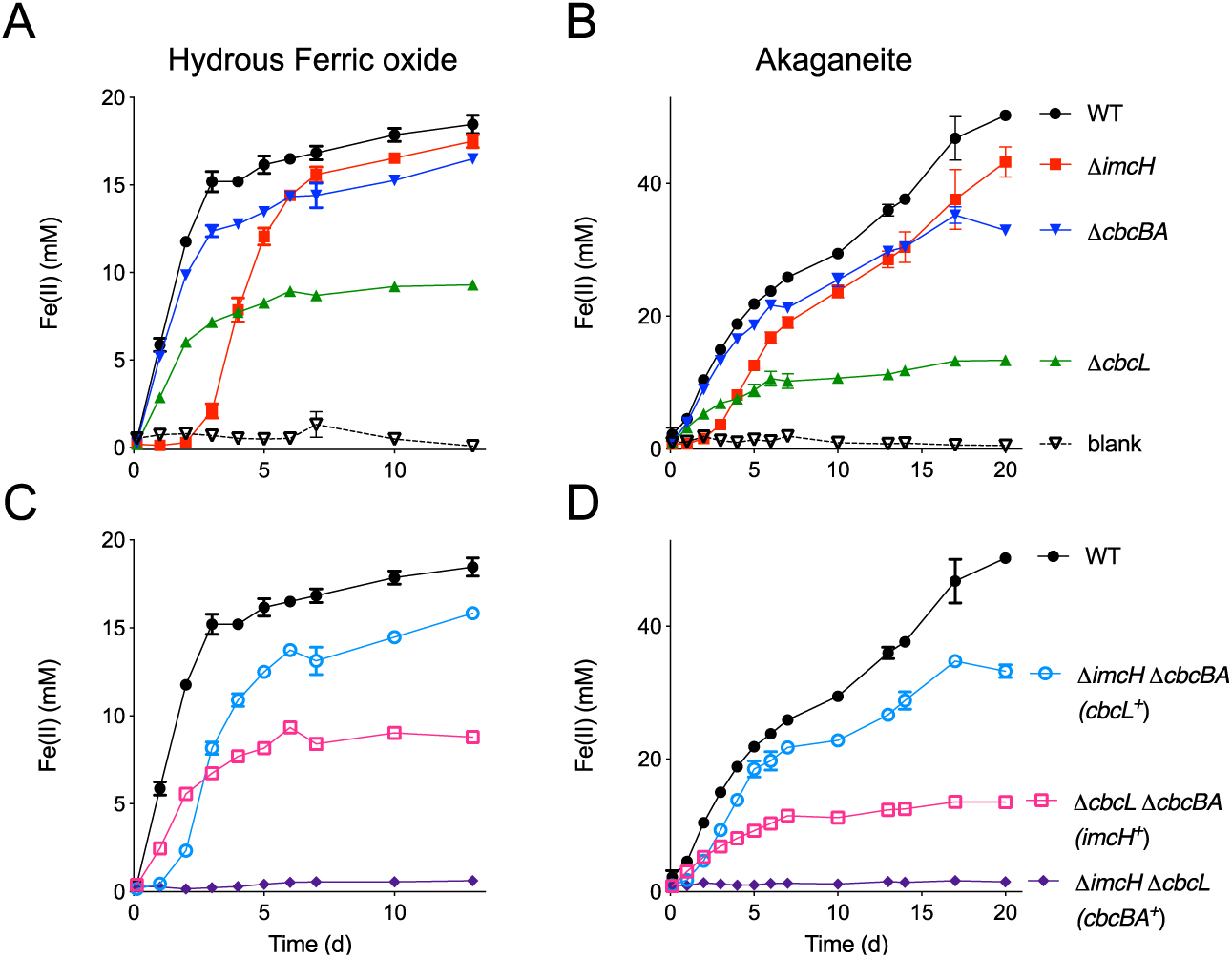
Complete reduction of Fe(III) oxides also requires *cbcBA*, regardless of Fe(III) (oxyhydr)oxide mineral preparation method. A and C. Reduction of hydrous ferric oxide, which has a formal redox potential lower than Fe(III) citrate, by single and double mutants lacking inner membrane cytochromes (Δ*imcH*, Δ*cbcL*, Δ*cbcBA*, Δ*imcH* Δ*cbcL*, Δ*cbcBA* Δ*cbcL*, and Δ*cbcBA* Δ*imcH*). A lag is observed by the Δ*imcH* mutant, but up to 95% of ferric oxide is eventually reduced. All mutants lacking Δ*cbcL* reduce 50% of Fe(III) oxide, and the Δ*cbcBA* mutant reduced only 90%. B and D. Reduction of akaganeite, which has a lower formal redox potential than hydrous ferric oxide, showed shorter lag for Δ*imcH*, and a larger defect for Δ*cbcL* mutants who could reduce only 26% of the Fe(III). Similarly, a larger defect was observed for the Δ*cbcBA* mutants in akaganeite. All experiments were conducted in triplicate, and the results are reported as mean values ± standard deviation.

The double deletion mutant Δ*imcH* Δ*cbcBA* (CbcL^+^) displayed the same lag as seen in Δ*imcH* but then also failed to reduce the last 10-15% of Fe(III) as seen for Δ*cbcBA* (Figure 6A, 6C). The double mutant Δ*cbcL* Δ*cbcBA* (ImcH^+^) ceased reduction similar to Δ*cbcL*, reducing 50% as much Fe(III) as WT (Figure 6C). Fe(III) reduction by double mutants aligned with the abilities of single mutants. Notably, even though concentrations of Fe(II) were much lower in hydrous ferric oxide incubations than in Fe(III) citrate, each cytochrome was necessary at the same phase of reduction, supporting the hypothesis that phenotypes were linked to the effective redox potential, not absolute Fe(III) or Fe(II) concentrations.

When a lower potential Fe(III) mineral (akaganeite) was used, the lag by Δ*imcH* was shorter (Figure 6B), consistent with less Fe(II) needing to accumulate to reduce redox potential and activate CbcL. Mutants lacking *cbcL* initiated growth, but only reduced 26% of Fe(III) compared to WT. Cells lacking *cbcBA* only reduced 65% of WT Fe(III) (Figure 6B). The extent of Fe(III) reduction by the double mutant Δ*cbcL* Δ*cbcBA* (ImcH^+^) was the same as Fe(III) reduction by Δ*cbcL*, and reduction by Δ*imcH* Δ*cbcBA* (CbcL^+^) was equivalent to reduction by the single mutant Δ*cbcBA* (Figure 6D).

These results across different electron acceptors, Fe(III) forms, and Fe(II) concentrations were consistent with ImcH, CbcL, and CbcBA each having a role at a different redox potential. In all cases, ImcH was essential when redox potential was above ∼0 V, CbcL was needed for reduction of moderately low potential acceptors (to about −0.2 V), and CbcBA was necessary for reduction closest to the thermodynamic limit. As lower potential electron acceptors such as akaganeite were used, CbcBA became more important for complete reduction.

While double mutants containing either *imcH* or *cbcL* demonstrated growth under at least one condition, double mutants containing only *cbcBA* failed to reduce Fe(III) (Figure 6C, 6D), and the same Δ*imcH* Δ*cbcL* mutant also failed to grow at any potential on electrodes (SI figure 5). The inability of cells containing only *cbcBA* to grow raised the possibility that CbcBA-dependent electron transfer conserves much less energy than when ImcH or CbcL is involved, possibly to the point where it cannot produce enough energy to support growth by *G. sulfurreducens* (SI figure 6). It also suggested that these are the only three options supporting Fe(III) reduction in this organism.

### Inner membrane cytochrome background affects growth yield

Similar to how oxygen-limited *E. coli* induces separate terminal oxidases with a lower proton pumping stoichiometry, an explanation for different quinone oxidoreductase-like genes in *Geobacter* could be generation of variable amounts of proton motive force in response to environmental conditions [70, 71]. Support for this hypothesis can be found in slower growth rates of electrode-reducing Δ*imcH* cells [31] and higher cell counts per mol Fe(II) in Δ*cbcL* cells [35]. However, strains in these prior experiments still contained *cbcBA*, which could have been contributing to phenotypes.

If cells containing ImcH translocate more protons than when a CbcL or CbcBA-dependent pathway is in use, then forcing cells to only use ImcH and not transition to use of the other pathways should increase ATP production and growth yield. In agreement with this prediction, *cbcL* deletion led to higher cell numbers at the end of Fe(III) reduction (Figure 7A). Cell counts increased further when both *cbcL* and *cbcBA* were deleted. When accounting for how much Fe(III) was reduced, these differences were even more pronounced (Figure 7B). Growth yield of Δ*cbcBA* increased 112 ± 25% compared to WT, yield of Δ*cbcL* increased 152 ± 32%, and yield of Δ*cbcBA* Δ*cbcL* (ImcH^+^) more than doubled, to 223 ± 59% (Figure 7B). This supported higher net ATP generation by ImcH-utilizing cells compared to those using CbcL or CbcBA.

**Figure 7.**
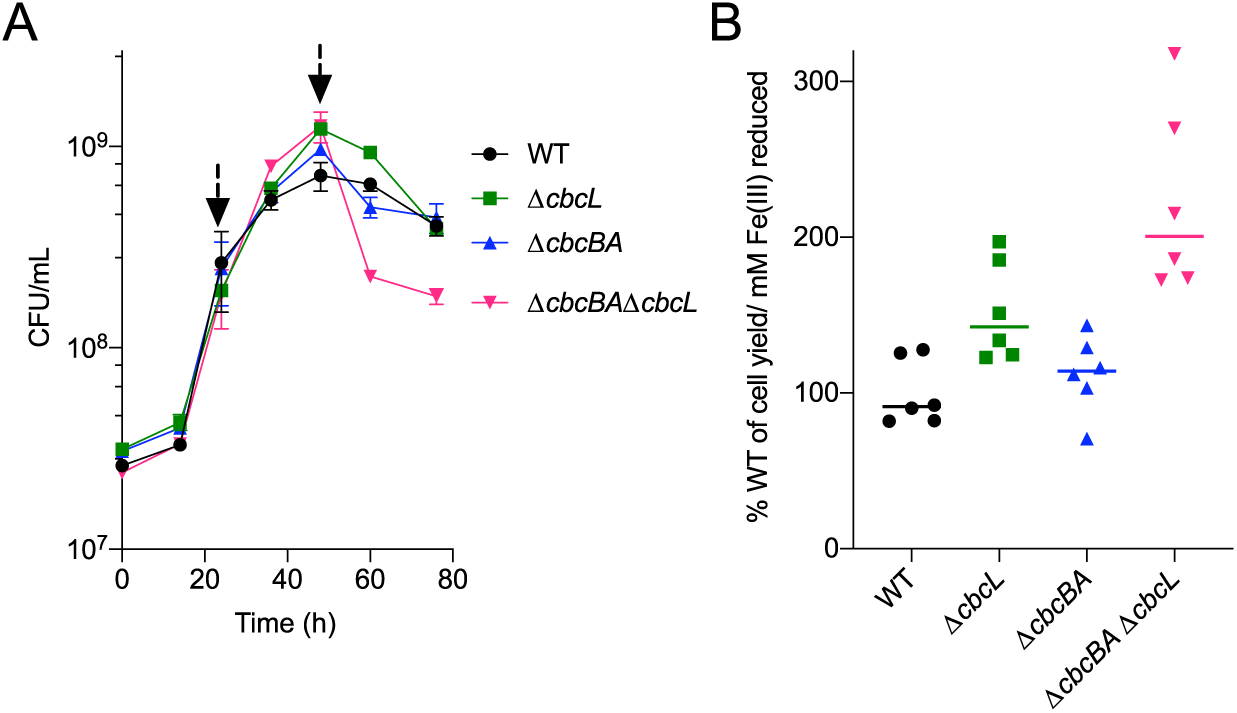
Cell yield (cells per mol Fe(III) reduced) increases in mutants lacking cbcL and cbcBA. A. Colony forming units (CFU/mL) measured from Fe(III) citrate-grown cultures. All mutants had similar initial growth rates, but mutants lacking *cbcBA* (Δ*cbcBA*, and Δ*cbcBA* Δ*cbcL*) showed a sharp decrease in viability compared to WT or Δ*cbcL*. Arrows represent the time interval used to calculate yield. B. Mutants lacking *cbcL*, *cbcBA*, or both had higher cellular yield per unit Fe(III) reduced. The ImcH-only strain (Δ*cbcBA* Δ*cbcL*) had the highest cellular yield, or 223% compared to WT, suggesting higher H+/e− compared to cultures where CbcL or CbcBA-dependent electron transfer initiated later in the growth curve. All experiments were conducted in triplicate (N=4), and data shown is represented as mean ± standard error of mean (SEM).

While CbcL and CbcBA negatively affected yield, data showed that these genes positively affect viability as Fe(III) became limiting. Near the end of Fe(III) reduction, viability of Δ*cbcBA* dropped over 50%, and Δ*cbcL* dropped by 68%. Cells lacking both *cbcL* and *cbcBA* had the worst survival, losing over 85% of cell viability within 24 h. A decrease in proton translocation stoichiometry would not only lower growth yield, but would also allow *G. sulfurreducens* to continue conserving energy as Fe(III) reduction becomes less favorable. Because we have been unable to demonstrate growth with extracellular electron acceptors by *cbcBA*-only strains, and CbcBA is necessary when less than 0.07 V/electron is available (7 kJ/e^−^), we hypothesize that CbcBA participates in an electron disposal route that primarily meets maintenance requirements when conditions are near thermodynamic limits.

## Discussion

Long before the isolation of metal-reducing bacteria, higher potential Mn(IV) in sediments was shown to be reduced before lower potential Fe(III) [5]. In this report, we provide a molecular explanation for how a microorganism can choose the most thermodynamically beneficial acceptor amid a collection of minerals that lie beyond the cell membrane. Our data supports a model where redox potential controls which of three different inner membrane respiratory pathways are used, removing the need to sense the solubility or chemistry of complex extracellular metal oxides in a changing environment.

Data from this study, combined with prior genetic observations [31, 32, 35] are consistent with *G. sulfurreducens* utilizing ImcH to achieve high growth rates and yields when redox potential is above −0.1 V. As redox potential decreases below this level, cells are increasingly dependent on CbcL, which lowers growth rate and yield but continues generating energy. As both of these cytochromes are constitutively expressed, this model predicts that CbcL should have a mechanism to prevent it from functioning at higher potentials. When redox potential approaches −0.2 V *vs.* SHE, induction of *cbcBA* provides a means for cells to respire if CbcL cannot function, and the energy available to the organism approaches zero (Figure 4). The fact that *cbcBA* is not expressed until it is needed is consistent with it supporting the lowest growth yields.

Although CbcBA and CbcL both have type I *b*-type diheme quinone oxidoreductase domains, they share no sequence homology, and have a different number of transmembrane helices predicted to coordinate their hemes. CbcB has four transmembrane domains, with 3 conserved histidines linked to *b*-heme coordination, based on alignments with characterized diheme proteins. While CbcA is a separate protein, a fourth histidine for binding a *b*-type heme appears to be located in its C-terminal domain. This pattern, where a *b*-type cytochrome is coordinated by a domain from a periplasmic enzyme is also seen in [NiFe] hydrogenases related to CbcB [72, 73].

CbcL has a different domain structure, with six transmembrane helices. One histidine capable of *b*-heme coordination is found in each of the first three transmembrane domains, but an additional two histidines arranged similar to those in formate dehydrogenase are in the fifth transmembrane domain [74]. The presence of five heme-coordinating residues could enable more than one *b*-heme binding configuration in CbcL, and provide a mechanism for preventing electron transfer until a key redox potential is reached. This hypothesis lacks precedent in other model systems and illustrates the need to biochemically characterize these putative quinone oxidoreductases.

Another feature of CbcBA is its consistent location in a regulated operon that is amongst one the most conserved cytochrome-encoding regions in *Geobacter*, occurring in 93 out of 96 *Geobacteraceae*, and 119 out of 134 Desulfuromonadales. Unlike *imcH* and *cbcL*, *cbcBA* is expressed only under low potential conditions (Figure 3). Our data here help explain studies that detected *cbcBA* expression in cells harvested after Fe(III) oxide reduction, but not higher-potential Mn(IV) oxide reduction [36]. Upregulation of *cbcBA* in electrode-grown biofilms is also consistent with *G. sulfurreducens* biofilms having low-potential regions farther from electrodes [25, 75, 76]. We predict that moderate *cbcBA* expression reported in electrode biofilms is an average of high expression in upper leaflets with low levels in the rest of the biofilm [76, 77]. Considering all of these studies, the radical change in *cbcBA* expression during growth with the same electron acceptor highlights a need to control or account for redox potential when cells are harvested for RNA extraction (SI figure 1).

Such fine-tuning of respiration is not found in all metal reducing organisms. *Shewanella oneidensis* uses one inner membrane quinone dehydrogenase, the tetraheme *c*-type cytochrome CymA [78] for reduction of acceptors that differ in redox potential by over 0.6 V, including fumarate, nitrate, DMSO, Fe(III), and Mn(IV) [79]. This may be explained by the fact that *Shewanella* partially oxidizes organic compounds to derive most of its ATP via substrate-level phosphorylation, and uses extracellular electron transfer primarily for electron disposal [80]. In contrast, *Geobacter* completely oxidizes substrates and requires chemiosmosis for ATP generation. Having multiple options for coupling electron flow to proton extrusion may allow *Geobacter* to utilize all available electrons and compete under such varied conditions as laboratory enrichments selecting for rapid growth, energy-limited aquifers selecting for persistence, and electrodes that create redox-stratified biofilms [75, 81, 82].

Nearly every important biological respiration can be easily identified by a highly conserved functional gene, such as *mcr* for methanogenesis, *dsr* for sulfate reduction, or *amo* for ammonia oxidation. Tools for molecular detection of metal-reducing bacteria are lacking, and prediction of extracellular electron transfer in uncultivated organisms is difficult, due to poor sequence similarity between multiheme cytochromes and poor conservation of cytochrome content between organisms [25, 37, 83, 84]. Unlike most *Geobacter c*-type cytochromes, the sequence of the *b*-type cytochrome CbcB is highly conserved, possibly because its donor (menaquinone) and acceptor (CbcA) remains more constant. This reduced rate of genetic drift allows CbcBA homologs near BccR-like regulators to be easily identified in other Deltaproteobacteria (such as metal-reducing *Anaeromyxobacter*) where the *b*-heme protein is typically annotated as ‘thiosulfate reductase’-like. Homologous *cbcBA* clusters annotated as hypothetical proteins are also present in metal-reducing genera such as the *Calditrichaeota* (*Caldithrix*) and *Bacteroidetes* (*Prolixibacter*, *Marinilabiliales*, *Labilibaculum*), making *cbcBA* a possible marker for extracellular electron transfer in more distant phyla. Based on the presence of *cbcBA* homologs in genomes from uncultivated organisms within the *Verrucomicrobia*, and a family of *cbcB-cbcA* gene fusions within *Chloroflexi* genomes, undiscovered organisms capable of extracellular respiration still remain buried deep within anoxic sediments and metagenomic bins.

## Acknowledgements

We thank the University of Minnesota Genomics Center for RNA sample processing and sequencing, and the University of Minnesota Supercomputing Institute for bioinformatic resources. This work was supported by the Office of Naval research grants N00014-16-1-2194, and N00014-18-1-2632.

---

**SI figure 1:**
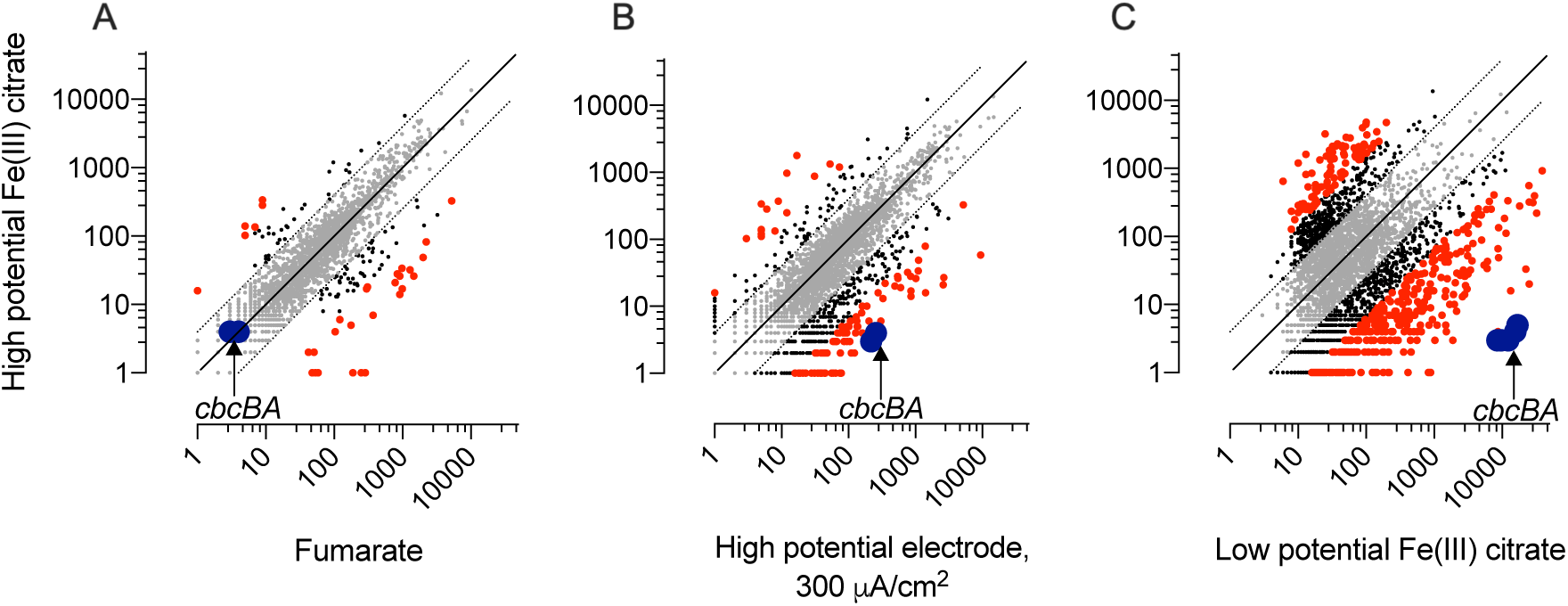
RNA-Seq of WT *G. sulfurreducens* grown with different electron acceptors compared to early exponential growth in Fe(III) citrate (30% reduced). A. Expression of *cbcBA* is barely detectable when both fumarate and high redox potential Fe(III) citrate are the electron acceptor. B. Expression of *cbcBA* increases in electrode-grown biofilms compared to fresh Fe(III) citrate, and C. Expression of *cbcBA* increases to over more than 10 000 RPKM (reads per kilobase mapped) in low potential Fe(III) citrate (70% reduced). Each comparison is the average of two biological replicates. Significant differences greater than 2-fold (black) or 4-fold (red) are highlighted. X and Y axes represent expression values.

**SI figure 2:**
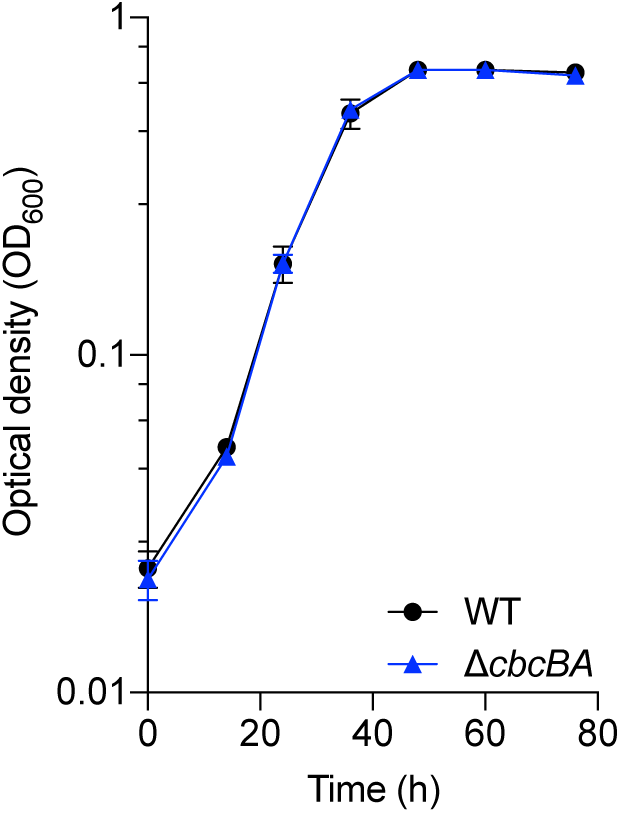
Growth of Δ*cbcBA* compared to WT in NB fumarate-acetate (NBFA) medium. Fully grown cultures of WT and Δ*cbcBA* in medium containing 40 mM fumarate and 20 mM acetate were inoculated 1:100 v/v, and optical density measured at 600 nm over time. Δ*cbcBA* did not show any growth defects as compared to WT when fumarate was the terminal electron acceptor (mean of three biological replicates, and error is reported as standard error of mean (SEM)).

**SI figure 3:**
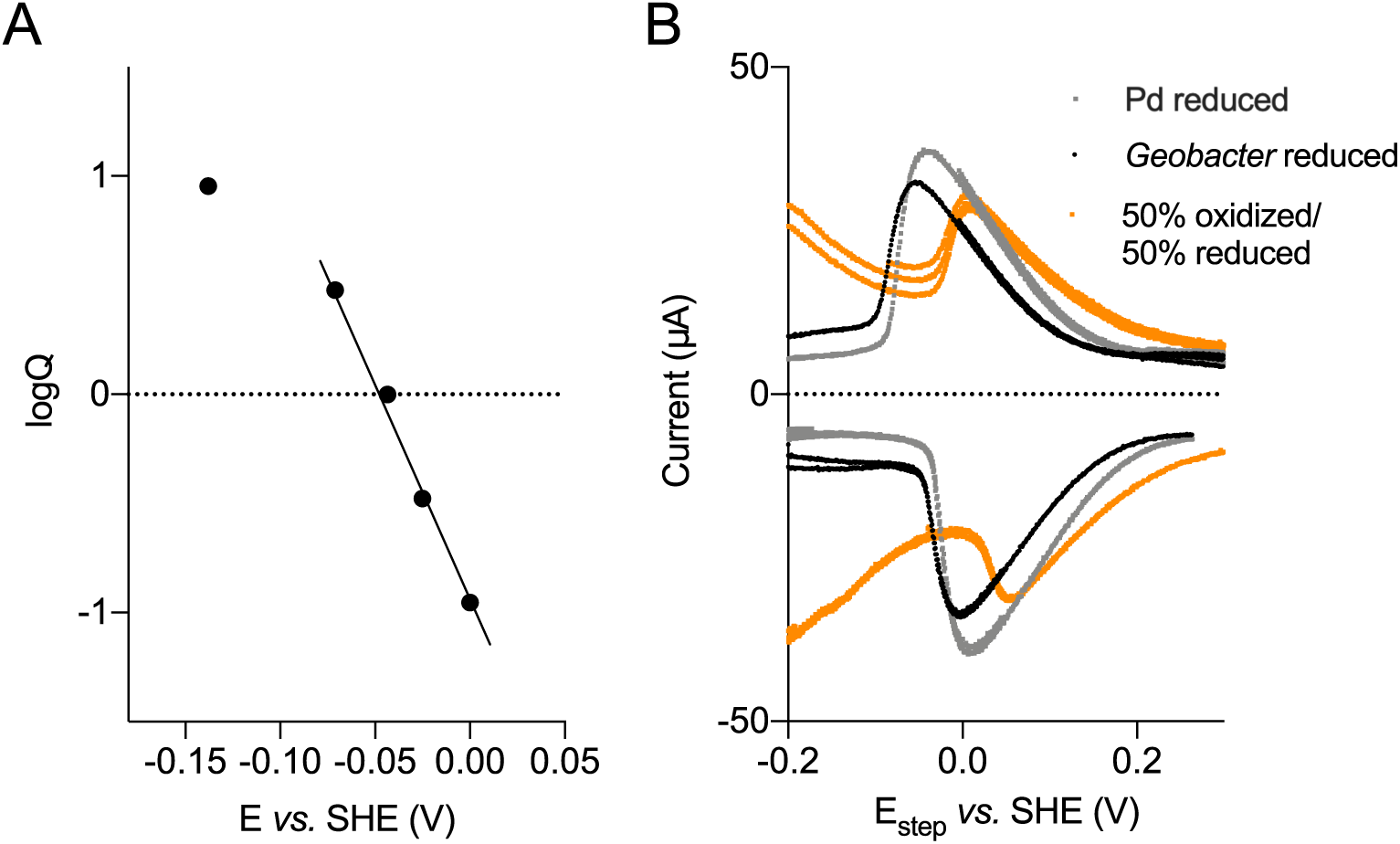
Calculating midpoint potential of Fe(III) citrate prepared in our laboratory. A. Calculation of midpoint potential of Fe(III) citrate using open circuit potential (OCP) method. Log(Fe(II)/Fe(III)) ratio of Fe(III) citrate follows Nernstian behavior only in a narrow potential window of about 0.1 V. Midpoint potential of Fe(III) citrate is 0.043 V *vs.* SHE. B. Comparing redox potential of Fe(III) citrate reduced by *G. sulfurreducens* or reduced by palladium (Pd) using differential pulse voltammetry method. These results show a much lower value of midpoint potential of Fe(III) citrate as previously reported [1] compared to the published values of +0.37 V *vs.* SHE [2].

**SI figure 4:**
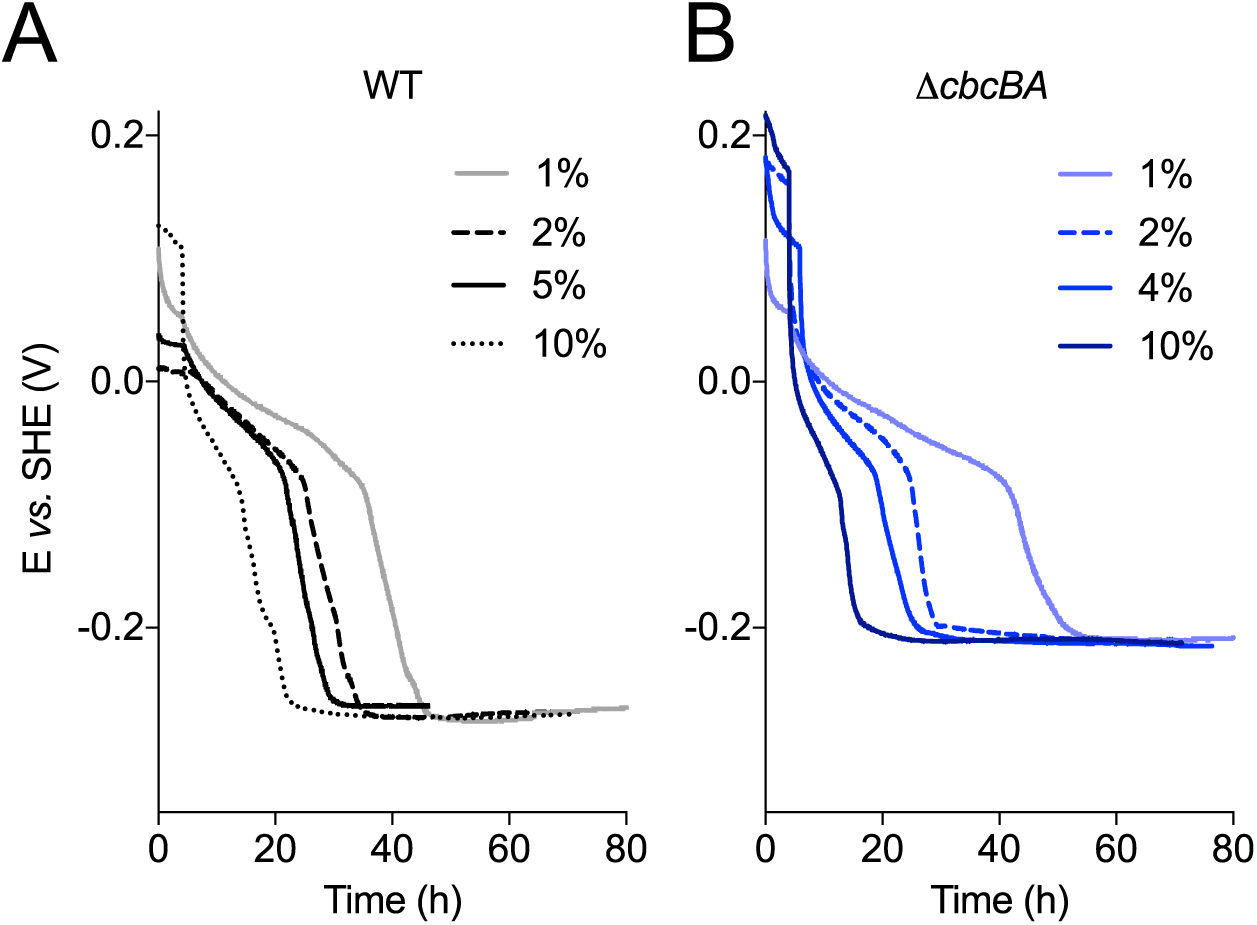
The extent of Fe(III) reduction does not change with the percentage of inoculation or the time of incubation. A. Redox potential of WT cultures reducing Fe(III) citrate when the percentage of inoculation was varied from 1% v/v to 10% v/v. The redox potential always stabilized at the same value. B. Redox potential of Δ*cbcBA* when inoculation was varied from 1% v/v to 10% v/v. Δ*cbcBA* always ceased reduction at a higher redox potential of *vs.* SHE, regardless of the percentage of inoculation. Redox potential was monitored for up to 80 h to test if Δ*cbcBA* cultures would lower the redox potential with longer incubation times. In similar experiments, the concentration of Fe(III) citrate was increased to 80 mM, and redox potential profiles followed similar trends, ending at the same final values [1].

**SI figure 5:**
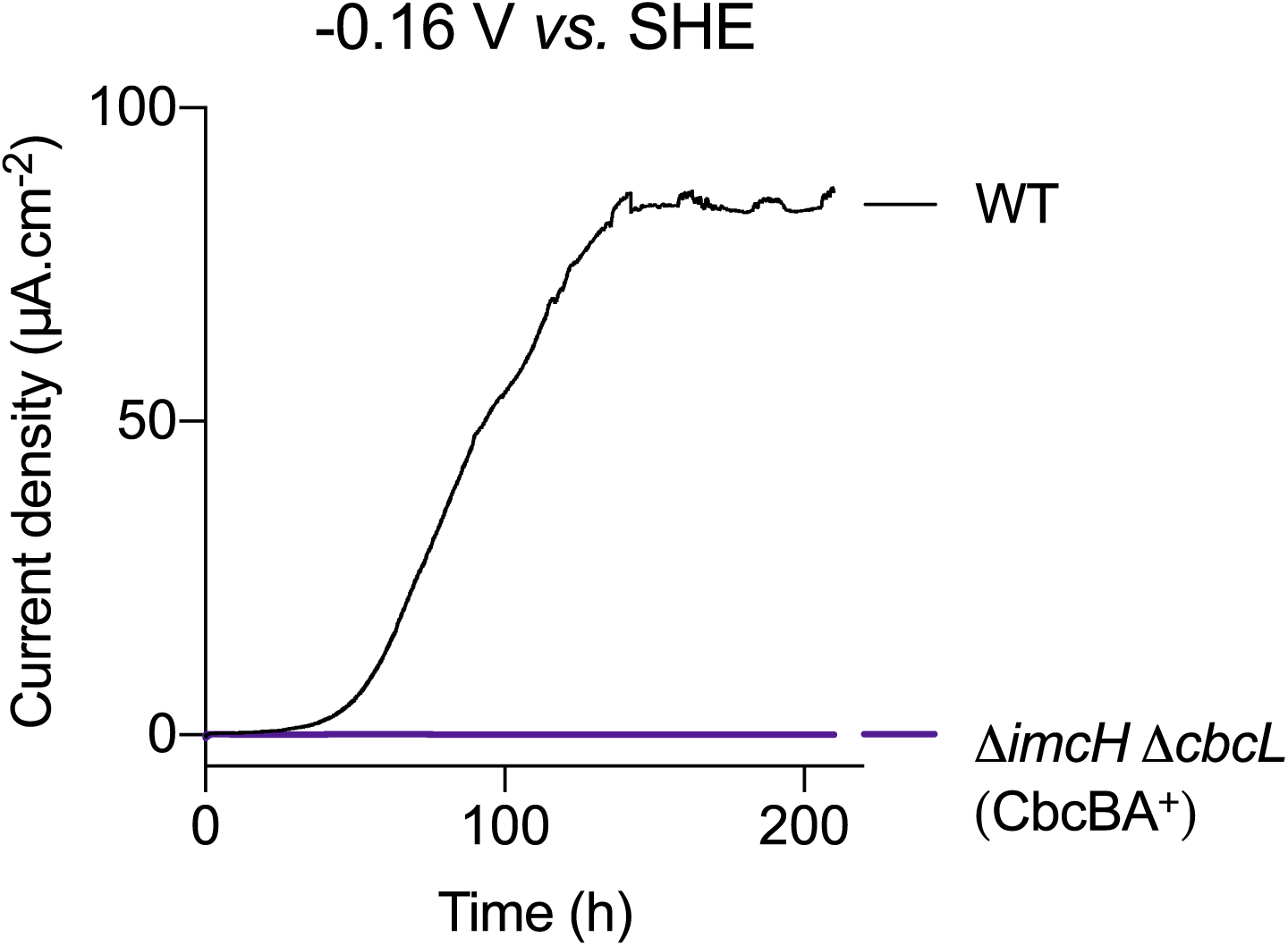
The Δ*imcH* Δ*cbcL* (CbcBA+) strain is unable to support growth on electrodes, even at lower potentials. When growth of Δ*imcH* Δ*cbcL* (CbcBA+) was tested with electrodes, shown here poised at −0.16 V *vs.* SHE as the terminal electron acceptor and acetate as the electron donor, CbcBA+ cells failed to produce any current as compared to WT. Experiments at −0.2 V also failed to produce current.

**SI figure 6:**
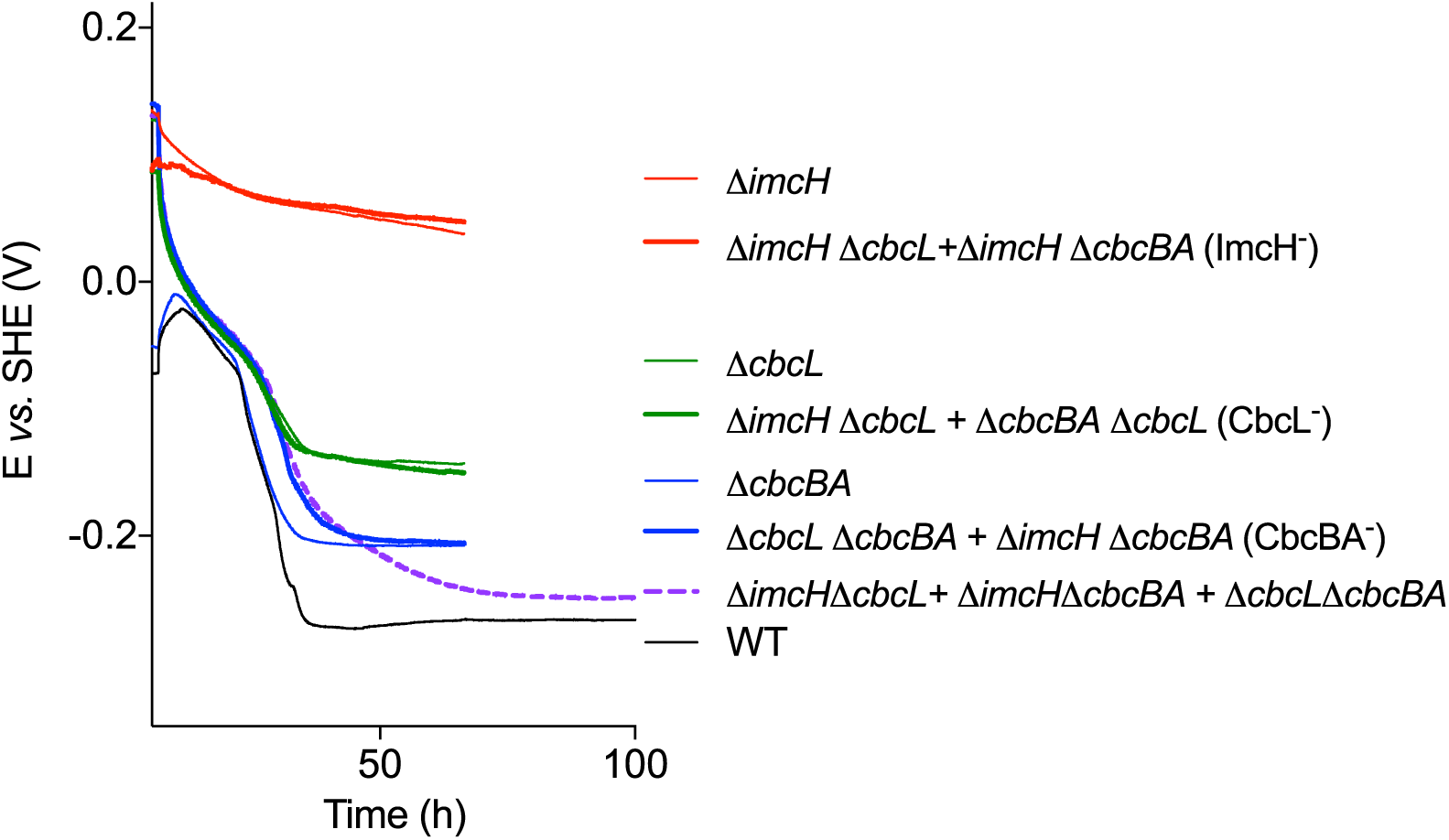
Combinations of mutants can ‘work together’, and produce the same final redox potential as mutants lacking the dominant inner membrane cytochrome. When reactors were inoculated with mixtures of double mutants, such as a mixture of cells lacking ImcH (Δ*imcH* Δ*cbcL* + Δ*imcH* Δ*cbcBA*), mixtures still failed to reduce any Fe(III) citrate, just as seen with Δ*imcH*. When mixtures of cells lacking CbcL (Δ*imcH* Δ*cbcL* + Δ*cbcL* Δ*cbcBA*) were inoculated, as one functional copy of *imcH* was present, together the mixture lowered redox potential to 0.15 V as observed with Δ*cbcL*. A combination of mutants, each lacking CbcBA (Δ*cbcL* Δ*cbcBA* + Δ*imcH* Δ*cbcBA*) together were able to reduce Fe(III) until redox potential reached 0.21 V *vs.* SHE, identical to the Δ*cbcBA* phenotype. was inoculated, redox potential went below the − .21 V *vs.* SHE threshold common to all When a mixture of all three double mutants (Δ*imcH* Δ*cbcL* + Δ*imcH* Δ*cbcBA* + Δ*cbcL* Δ*cbcBA*) Δ*cbcBA* mutants, but showed a slower rate of redox potential drop near the end of the incubation. A hypothesis for this behavior is the Δ*imcH* Δ*cbcL* cells which should induce *cbcBA* and finish reduction were unable to generate enough energy to fully produce enough *cbcBA*. Representative curves are shown from experiments (N=2) performed in duplicates.

## References

1. [1] Haddock, BA, Jones, CW (1977). Bacterial respiration. Bacteriol Rev 41(1): 47–99.

2. [2] Fischer, WR (1987). Standard potentials (Eo) of iron(III) oxides under reducing conditions. J Plant Nutr Soil Sci 150(5): 286–289.

3. [3] Thauer, RK, Jungermann, K, Decker, K (1977). Energy conservation in chemotrophic anaerobic bacteria. Bacteriol Rev 41(1): 100–180.

4. [4] ZoBell, CE (1946). Studies on redox potential of marine sediments. Am Assoc Pet Geol Bull 30(4): 477–513.

5. [5] Takai, Y, Kamura, T (1966). The mechanism of reduction in waterlogged paddy soil. Folia Microbiol 11(4): 304–313.

6. [6] Bethke, CM, Sanford, RA, Kirk, MF, Jin, Q, Flynn, TM (2011). The thermodynamic ladder in geomicrobiology. Am J Sci 311(3): 183–210.

7. [7] Simon, J, van Spanning, RJM, Richardson, DJ (2008). The organisation of proton motive and non-proton motive redox loops in prokaryotic respiratory systems. Biochim Biophys Acta - Bioenergetics 1777(12): 1480–1490.

8. [8] Unden, G, Dünnwald, P (2008). The aerobic and anaerobic respiratory chain of *Escherichia coli* and *Salmonella enterica*: enzymes and energetics. EcoSal Plus 3(1).

9. [9] Buckel, W, Thauer, RK (2018). Flavin-based electron bifurcation, a new mechanism of biological energy coupling. Chem Rev 118(7): 3862–3886.

10. [10] Lovley, DR (1991). Dissimilatory Fe(III) and Mn(IV) reduction. Microbiol Rev 55(2): 259–287.

11. [11] Nealson, KH, Saffarini, D (1994). Iron and manganese in anaerobic respiration: environmental significance, physiology, and regulation. Annu Rev Microbiol 48: 311–343.

12. [12] Thamdrup, B (2000). Bacterial manganese and iron reduction in aquatic sediments. In Schink, B (Ed.) *Advances in Microbial Ecology*, *Springer US, Boston*, MA 41–84.

13. [13] Straub, KL, Benz, M, Schink, B (2001). Iron metabolism in anoxic environments at near neutral pH. FEMS Microbiol Ecol 34(3): 181–186.

14. [14] Cornell, RM, Schwertmann, U (2006). The iron oxides: structure, properties, reactions, occurrences and uses. Wiley-VCH.

15. [15] Navrotsky, A, Mazeina, L, Majzlan, J (2008). Size-driven structural and thermodynamic complexity in iron oxides. Science 319(5870): 1635–1638.

16. [16] Lovley, DR, Phillips, EJP (1988). Manganese inhibition of microbial iron reduction in anaerobic sediments. Geomicrobiol J 6(3–4): 145–155.

17. [17] Lovley, DR, Phillips, EJP (1988). Novel mode of microbial energy-metabolism - organic-carbon oxidation coupled to dissimilatory reduction of iron or manganese. Appl Environ Microbiol 54(6): 1472–1480.

18. [18] Lovley, DR, Giovannoni, SJ, White, DC, Champine, JE, Phillips, EJP, Gorby, YA, et al. (1993). *Geobacter metallireducens* gen. nov. sp. nov., a microorganism capable of coupling the complete oxidation of organic compounds to the reduction of iron and other metals. Arch Microbiol 159(4): 336–344.

19. [19] Caccavo, F, Lonergan, DJ, Lovley, DR, Davis, M, Stolz, JF, McInerney, MJ (1994). *Geobacter sulfurreducens* sp. nov., a hydrogen- and acetate-oxidizing dissimilatory metal-reducing microorganism. Appl Environ Microbiol 60(10): 3752–3759.

20. [20] Anderson, RT, Vrionis, HA, Ortiz-Bernad, I, Resch, CT, Long, PE, Dayvault, R, et al. (2003). Stimulating the *in situ* activity of *Geobacter* species to remove uranium from the groundwater of a uranium-contaminated aquifer. Appl Environ Microbiol 69(10): 5884–5891.

21. [21] Rotaru, AE, Shrestha, PM, Liu, F, Markovaite, B, Chen, S, Nevin, KP, et al. (2014). Direct interspecies electron transfer between *Geobacter metallireducens* and *Methanosarcina barkeri*. Appl Environ Microbiol 80(15): 4599–4605.

22. [22] Bond, DR, Lovley, DR (2003). Electricity production by *Geobacter sulfurreducens* attached to electrodes. Appl Environ Microbiol 69(3): 1548–1555.

23. [23] Lloyd, JR, Leang, C, Hodges Myerson, AL, Coppi, MV, Cuifo, S, Methe, B, et al. (2003). Biochemical and genetic characterization of PpcA, a periplasmic *c*-type cytochrome in *Geobacter sulfurreducens*. Biochem J 369(Pt 1): 153–161.

24. [24] Shelobolina, ES, Coppi, MV, Korenevsky, AA, DiDonato, LN, Sullivan, SA, Konishi, H, et al. (2007). Importance of *c*-type cytochromes for U(VI) reduction by *Geobacter sulfurreducens*. BMC Microbiol 7: 16.

25. [25] Otero, FJ, Chan, CH, Bond, DR (2018). Identification of different putative outer membrane electron conduits necessary for Fe(III) citrate, Fe(III) oxide, Mn(IV) oxide, or electrode reduction by *Geobacter sulfurreducens*. J Bacteriol 200(19).

26. [26] Wang, F, Gu, Y, O’Brien, JP, Yi, SM, Yalcin, SE, Srikanth, V, et al. (2019). Structure of microbial nanowires reveals stacked hemes that transport electrons over micrometers. Cell 177(2): 361–369.e10.

27. [27] Yalcin, SE, O’Brien, JP, Gu, Y, Reiss, K, Yi, SM, Jain, R, et al. (2020). Electric field stimulates production of highly conductive microbial OmcZ nanowires. Nat Chem Bio 16(10): 1136–1142.

28. [28] Reguera, G, McCarthy, KD, Mehta, T, Nicoll, JS, Tuominen, MT, Lovley, DR (2005). Extracellular electron transfer via microbial nanowires. Nature 435(7045): 1098–1101.

29. [29] Jahan, MI, Tobe, R, Mihara, H (2018). Characterization of a novel porin-like protein, ExtI, from *Geobacter sulfurreducens* and its implication in the reduction of selenite and tellurite. Int J Mol Sci 19(3).

30. [30] Chan, CH, Levar, CE, Jiménez-Otero, F, Bond, DR (2017). Genome scale mutational analysis of *Geobacter sulfurreducens* reveals distinct molecular mechanisms for respiration and sensing of poised electrodes versus Fe(III) oxides. J Bacteriol 199(19).

31. [31] Levar, CE, Chan, CH, Mehta-Kolte, MG, Bond, DR (2014). An inner membrane cytochrome required only for reduction of high redox potential extracellular electron acceptors. mBio 5(6): e02 034–14.

32. [32] Zacharoff, L, Chan, CH, Bond, DR (2016). Reduction of low potential electron acceptors requires the CbcL inner membrane cytochrome of *Geobacter sulfurreducens*. Bioelectrochemistry 107: 7–13.

33. [33] Finkelstein, DA, Tender, LM, Zeikus, JG (2006). Effect of electrode potential on electrode-reducing microbiota. Environ Sci Technol 40(22): 6990–6995.

34. [34] Logan, BE, Hamelers, B, Rozendal, R, Schröder, U, Keller, J, Freguia, S, et al. (2006). Microbial fuel cells: methodology and technology. Environ Sci Technol 40(17): 5181–5192.

35. [35] Levar, CE, Hoffman, CL, Dunshee, AJ, Toner, BM, Bond, DR (2017). Redox potential as a master variable controlling pathways of metal reduction by *Geobacter sulfurreducens*. ISME J 11(3): 741–752.

36. Aklujkar, M, Coppi, MV, Leang, C, Kim, BC, Chavan, MA, Perpetua, LA, et al. (2013). Proteins involved in electron transfer to Fe(III) and Mn(IV) oxides by Geobacter sulfurreducens and Geobacter uraniireducens. Microbiology 159(Pt 3): 515–535.

37. [37] Butler, JE, Young, ND, Lovley, DR (2010). Evolution of electron transfer out of the cell: comparative genomics of six *Geobacter* genomes. BMC Genomics 11: 40.

38. [38] Chan, CH, Levar, CE, Zacharoff, L, Badalamenti, JP, Bond, DR (2015). Scarless genome editing and stable inducible expression vectors for *Geobacter sulfurreducens*. Appl Environ Microbiol 81(20): 7178–7186.

39. [39] Hallberg, ZF, Chan, CH, Wright, TA, Kranzusch, PJ, Doxzen, KW, Park, JJ, et al. (2019). Structure and mechanism of a Hypr GGDEF enzyme that activates cGAMP signaling to control extracellular metal respiration. eLife 8: e43 959.

40. [40] Ferriéres, L, Hémery, G, Nham, T, Guérout, AM, Mazel, D, Beloin, C, et al. (2010). Silent mischief: Bacteriophage Mu insertions contaminate products of *Escherichia coli* random mutagenesis performed using suicidal transposon delivery plasmids mobilized by broad-host-range RP4 conjugative machinery. J Bacteriol 192(24): 6418–6427.

41. [41] Peters, JM, Koo, BM, Patino, R, Heussler, GE, Hearne, CC, Qu, J, et al. (2019). Enabling genetic analysis of diverse bacteria with Mobile-CRISPRi. Nat Microbiol 4(2): 244–250.

42. [42] Marsili, E, Rollefson, JB, Baron, DB, Hozalski, RM, Bond, DR (2008). Microbial biofilm voltammetry: direct electrochemical characterization of catalytic electrode-attached biofilms. Appl Environ Microbiol 74(23): 7329–7337.

43. [43] Joshi, K, Kane, AL, Kotloski, NJ, Gralnick, JA, Bond, DR (2019). Preventing hydrogen disposal increases electrode utilization efficiency by *Shewanella oneidensis*. Front Energy Res 7(95).

44. [44] Lovley, DR, Phillips, EJP (1987). Rapid assay for microbially reducible ferric iron in aquatic sediments. Appl Environ Microbiol 53(7): 1536–1540.

45. [45] Langmead, B, Salzberg, SL (2012). Fast gapped-read alignment with Bowtie 2. Nat Methods 9(4): 357–359.

46. [46] McClure, R, Balasubramanian, D, Sun, Y, Bobrovskyy, M, Sumby, P, Genco, CA, et al. (2013). Computational analysis of bacterial RNA-Seq data. Nucleic Acids Res 41(14): e140.

47. [47] Herigstad, B, Hamilton, M, Heersink, J (2001). How to optimize the drop plate method for enumerating bacteria. J Microbiol Methods 44(2): 121–129.

48. Methé, BA, Nelson, KE, Eisen, JA, Paulsen, IT, Nelson, W, Heidelberg, JF, et al. (2003). Genome of *Geobacter sulfurreducens*: Metal reduction in subsurface environments. Science 302(5652): 1967–1969.

49. [49] Qu, Y, Brown, P, Barbe, JF, Puljic, M, Merino, E, Adkins, RM, et al. (2009). GSEL version 2, an online genome-wide query system of operon organization and regulatory sequence elements of *Geobacter sulfurreducens*. OMICS 13(5): 439– 449.

50. [50] Krushkal, J, Leang, C, Barbe, JF, Qu, Y, Yan, B, Puljic, M, et al. (2009). Diversity of promoter elements in a *Geobacter sulfurreducens* mutant adapted to disruption in electron transfer. Funct Integr Genomics 9(1): 15–25.

51. [51] Yan, B, Núñez, C, Ueki, T, Esteve-Núñez, A, Puljic, M, Adkins, RM, et al. (2006). Computational prediction of RpoS and RpoD regulatory sites in *Geobacter sulfurreducens* using sequence and gene expression information. Gene 384: 73–95.

52. [52] Holmes, DE, Chaudhuri, SK, Nevin, KP, Mehta, T, Methé, BA, Liu, A, et al. (2006). Microarray and genetic analysis of electron transfer to electrodes in *Geobacter sulfurreducens*. Environ Microbiol 8(10): 1805–1815.

53. [53] Krogh, A, Larsson, B, von Heijne, G, Sonnhammer, EL (2001). Predicting transmembrane protein topology with a hidden Markov model: application to complete genomes. J Mol Biol 305(3): 567–580.

54. [54] Yu, NY, Wagner, JR, Laird, MR, Melli, G, Rey, S, Lo, R*, et al.* (2010). PSORTb 3.0: Improved protein subcellular localization prediction with refined localization subcategories and predictive capabilities for all prokaryotes. Bioinformatics 26(13): 1608–1615.

55. [55] Ding, YHR, Hixson, KK, Giometti, CS, Stanley, A, Esteve-Núñez, A, Khare, T, et al. (2006). The proteome of dissimilatory metal-reducing microorganism *Geobacter sulfurreducens* under various growth conditions. Biochim Biophys Acta-Proteins and Proteomics 1764(7): 1198–1206.

56. [56] Leang, C, Krushkal, J, Ueki, T, Puljic, M, Sun, J, Juárez, K, et al. (2009). Genome-wide analysis of the RpoN regulon in *Geobacter sulfurreducens*. BMC Genomics 10: 331.

57. [57] Blum, M, Chang, HY, Chuguransky, S, Grego, T, Kandasaamy, S, Mitchell, A, et al. (2021). The InterPro protein families and domains database: 20 years on. Nucleic Acids Res 49(D1): D344–D354.

58. [58] Brasca, M, Morandi, S, Lodi, R, Tamburini, A (2007). Redox potential to discriminate among species of lactic acid bacteria. J Appl Microbiol 103(5): 1516– 1524.

59. [59] Vukosav, P, Mlakar, M, Tomišić, V (2012). Revision of iron(III)–citrate speciation in aqueous solution. Voltammetric and spectrophotometric studies. Anal Chim Acta 745: 85–91.

60. [60] Adam, FI, Bounds, PL, Kissner, R, Koppenol, WH (2015). Redox properties and activity of iron–citrate complexes: Evidence for redox cycling. Chem Res Toxicol 28(4): 604–614.

61. [61] Königsberger, LC, Königsberger, E, May, PM, Hefter, GT (2000). Complexation of iron(III) and iron(II) by citrate. Implications for iron speciation in blood plasma. J Inorg Biochem 78(3): 175–184.

62. [62] Baerends, RJ, Smits, WK, de Jong, A, Hamoen, LW, Kok, J, Kuipers, OP (2004). Genome2D: a visualization tool for the rapid analysis of bacterial transcriptome data. Genome Biol 5(5): R37.

63. [63] Ueki, T, Lovley, DR (2010). Genome-wide gene regulation of biosynthesis and energy generation by a novel transcriptional repressor in *Geobacter* species. Nucleic Acids Res 38(3): 810–821.

64. [64] Richter, H, Nevin, KP, Jia, H, Lowy, DA, Lovley, DR, Tender, LM (2009). Cyclic voltammetry of biofilms of wild type and mutant *Geobacter sulfurreducens* on fuel cell anodes indicates possible roles of OmcB, OmcZ, type IV pili, and protons in extracellular electron transfer. Energy Environ Sci 2(5): 506–516.

65. [65] Yoho, RA, Popat, SC, Torres, CI (2014). Dynamic potential-dependent electron transport pathway shifts in anode biofilms of *Geobacter sulfurreducens*. ChemSusChem 7(12): 3413–3419.

66. [66] Marsili, E, Sun, J, Bond, D (2010). Voltammetry and growth physiology of *Geobacter sulfurreducens* biofilms as a function of growth stage and imposed electrode potential. Electroanalysis 22(7-8): 865–874.

67. [67] Bonneville, S, Behrends, T, Van Cappellen, P (2009). Solubility and dissimilatory reduction kinetics of iron(III) oxyhydroxides: A linear free energy relationship. Geochim Cosmochim Acta 73(18): 5273–5282.

68. [68] Majzlan, J, Navrotsky, A, Schwertmann, U (2004). Thermodynamics of iron oxides: Part III. Enthalpies of formation and stability of ferrihydrite (Fe(OH)3), schwertmannite (FeO(OH)3/4(SO4)1/8), and ɛ-Fe2O3. Geochim Cosmochim Acta 713 68(5): 1049–1059.

69. [69] Majzlan, J (2012). Minerals and aqueous species of iron and manganese as reactants and products of microbial metal respiration. In Gescher, J, Kappler, A (Eds.) Microbial Metal Respiration, Springer Berlin Heidelberg, Berlin, Heidelberg 1– 28.

70. [70] Ingledew, WJ, Poole, RK (1984). The respiratory chains of *Escherichia coli*. Microbiol Rev 48(3): 222–271.

71. [71] Bekker, M, Vries, Sd, Beek, AT, Hellingwerf, KJ, Mattos, MJTD (2009). Respiration of *Escherichia coli* can be fully uncoupled via the nonelectrogenic terminal cytochrome *bd*-II oxidase. J Bacteriol 191(17): 5510–5517.

72. [72] Gross, R, Simon, J, Lancaster, CRD, Kröger, A (1998). Identification of histidine residues in *Wolinella succinogenes* hydrogenase that are essential for menaquinone reduction by H2. Mol Microbiol 30(3): 639–646.

73. [73] Gross, R, Pisa, R, Sänger, M, Lancaster, CRD, Simon, J (2004). Characterization of the menaquinone reduction site in the diheme cytochrome *b* membrane anchor of *Wolinella succinogenes* NiFe-hydrogenase. J Biol Chem 279(1): 274–281.

74. [74] Jormakka, M, Törnroth, S, Byrne, B, Iwata, S (2002). Molecular basis of proton motive force generation: structure of formate dehydrogenase-N. Science 295(5561): 1863–1868.

75. [75] Chadwick, GL, Otero, FJ, Gralnick, JA, Bond, DR, Orphan, VJ (2019). NanoSIMS imaging reveals metabolic stratification within current-producing biofilms. Proc Natl Acad Sci U S A 116(41): 20 716–20 724.

76. [76] Krige, A, Ramser, K, Sjöblom, M, Christakopoulos, P, Rova, U (2020). A new approach for evaluating electron transfer dynamics by using *in situ* resonance raman microscopy and chronoamperometry in conjunction with a dynamic model. Appl Environ Microbiol 86(20): 320–327.

77. [77] Lebedev, N, Strycharz-Glaven, SM, Tender, LM (2014). Spatially resolved confocal resonant Raman microscopic analysis of anode-grown *Geobacter sulfurreducens* biofilms. Chemphyschem 15(2): 320–327.

78. [78] Myers, CR, Myers, JM (1997). Cloning and sequence of *cymA*, a gene encoding a tetraheme cytochrome *c* required for reduction of iron(III), fumarate, and nitrate by *Shewanella putrefaciens* MR-1. J Bacteriol 179(4): 1143–1152.

79. [79] Marritt, SJ, Lowe, TG, Bye, J, McMillan, DGG, Shi, L, Fredrickson, J, et al. (2012). A functional description of CymA, an electron-transfer hub supporting anaerobic respiratory flexibility in *Shewanella*. Biochem J 444(3): 465–474.

80. [80] Hunt, KA, Flynn, JM, Naranjo, B, Shikhare, ID, Gralnick, JA (2010). Substrate-level phosphorylation is the primary source of energy conservation during anaerobic respiration of *Shewanella oneidensis* strain MR-1. J Bacteriol 192(13): 3345–3351.

81. [81] Snider, RM, Strycharz-Glaven, SM, Tsoi, SD, Erickson, JS, Tender, LM (2012). Long-range electron transport in *Geobacter sulfurreducens* biofilms is redox gradient-driven. Proc Natl Acad Sci U S A 109(38): 15 467–15 472.

82. [82] He, X, Chadwick, G, Otero, FJ, Orphan, V, Meile, C (2021). Spatially resolved electron transport through anode-respiring *Geobacter sulfurreducens* biofilms: controls and constraints. ChemElectroChem (in press).

83. [83] Meyer, TE, Kamen, MD (1982). New perspectives on *c*-type cytochromes. Adv Protein Chem 35: 105–212.

84. [84] Scott Mathews, F (1985). The structure, function and evolution of cytochromes. Prog Biophys Mol Biol 45(1): 1–56.

85. [85] Choi, KH, Mima, T, Casart, Y, Rholl, D, Kumar, A, Beacham, IR, et al. (2008). Genetic tools for select-agent-compliant manipulation of *Burkholderia pseudomallei*. Appl Environ Microbiol 74(4): 1064–1075.

